# Hormonal dynamics reveal a stimulatory role for secretoneurin in zebrafish ovulation

**DOI:** 10.1101/2024.07.03.601258

**Authors:** Di Peng, Chunyu Lu, Victoria Spadacini, Kimberly Mitchell, Yongjun Tan, Dapeng Zhang, Berta Levavi-Sivan, Wei Hu, Vance L. Trudeau

## Abstract

Surge release of luteinizing hormone (Lh) from the pituitary is essential for fertility as it triggers ovulation. While secretoneurin (SN) is a phylogenetically conserved secretogranin-2 derived peptide that stimulates Lh, its role in ovulation has not been established. To directly compare periovulatory changes in the classical hormones to the emerging reproductive neuropeptides SNa and SNb, simultaneous mass spectrometry measurement of 9 peptides and 5 steroids was conducted in female zebrafish. Regression analysis indicated that levels of SNa1-34 in the brain peaked when type 3 gonadotropin-releasing hormone (Gnrh3) increased (R^2^=0.71) at the time of the Lh surge, 3.5 h before ovulation. In contrast, levels of the naturally occurring derivative SNa1-14 were highest at ovulation, while SNb1-31 was invariable. The bioactivities of SNa1-34 and SNa1-14 were investigated. After injection of SNa1-34 in females that had been isolated from males, 61% (11/18) ovulated within 6 h, which was like the effects of the Lh analog human chorionic gonadotropin (hCG) (72%; 13/18 females). SNa1-34 could induce ovulation by stimulating time-dependent expression of *gnrh3* in the brain, *chorionic gonadotropin alpha (cga), luteinizing hormone b (lhb)* subunit in pituitary, and the *nuclear progesterone receptor (npr)* in ovaries. In contrast, SNa1-14 exhibited far fewer effects on gene expression and did not induce ovulation. Our results support the proposal that SN is a reproductive hormone.

**Significance statement:** Secretogranin-2 is a secretory granule protein that is enzymatically processed to the bioactive neuropeptide secretoneurin. It is produced in hypothalamic neurons and anterior pituitary cells, and we provide *in vivo* evidence that secretoneurin plays an important role to induce ovulation. Secretoneurin levels in the brain increase concomitantly with gonadotropin-releasing hormone prior to ovulation in normal zebrafish. Injection of a synthetic secretoneurin peptide increased expression of reproduction-related genes at all levels of the hypothalamic-pituitary-ovarian axis. Secretoneurin stimulates ovulation in otherwise anovulatory females. Together these data place the evolutionarily conserved secretoneurin amongst other critical neuroendocrine regulators. Secretoneurin or synthetic agonists could be used to improve breeding in fish species, or to potentially help with infertility treatments

## Introduction

The coordinated synthesis and release of hormones of the hypothalamus-pituitary-gonadal (HPG) axis control timed ovulation in females and sperm release in males, thus determining fertility in vertebrate species. A central concept is that gonadotropin-releasing hormone 1 (Gnrh1) neurons perform critical roles in mammalian reproduction. Hypothalamic neurons projecting to the median eminence release Gnrh1 into the portal blood system, activating Gnrh receptors, and stimulate the synthesis and secretion of the gonadotropins follicle stimulating hormone (Fsh) and luteinizing hormone (Lh) from the anterior pituitary (1). These gonadotropins in turn activate their receptors to regulate ovarian steroid production, oocyte development, key morphological changes, and ultimately ovulation (2). The gonadal steroids released into the blood feedback both positively and negatively on neurons and pituitary cells to finely control gonadotropin secretion, driving the ovulatory process.

Many other neuropeptides produced in the hypothalamus are also involved, and the kisspeptins have emerged as key players in mammals (3, 4). Kisspeptin (Kiss) directly controls Gnrh1 neurons to regulate Lh release (5). Hypothalamic arcuate neurons co-expressing Kiss, neurokinin B (Nkb), and dynorphin A serve as a Gnrh pulse generator that drives the preovulatory Lh surge. These Kisspeptin/Neurokinin B/Dynorphin neurons are essential to normal reproduction in mice (6).

Critically, data in emerging models for reproductive physiology such as zebrafish and medaka now challenge this concept of essentiality of Gnrh and Kiss (7, 8). Unlike mammals, the neural mechanisms that trigger a preovulatory Lh surge after folliculogenesis remain unclear in fish and other non-mammalian vertebrates. In teleost fish the median eminence was lost over evolutionary time. Therefore a multitude of hypothalamic neurons, including those that produce the fish Gnrh and Kiss peptides, terminate near, or directly innervate, the distinct gonadotroph cells releasing Lh or Fsh (9), thus conferring control over them without a tetrapod-like median eminence-portal blood system. In zebrafish, Gnrh2/Gnrh3 single or double mutants (10, 11) and those with chemogenic depletion of Gnrh3 are fertile (12). Medaka harboring mutations in *gnrh1* have well-developed ovaries but fail to ovulate. In these female medaka, mutation of *gnrh1* did not affect *fshb* and had only a minor suppressive effect on *lhb* whereby gonadal development was maintained. Intriguingly, *gnrh1* mutant male medaka remained fertile (13). Equally surprising is that numerous zebrafish *kiss* or *kissr* mutant lines reproduce normally (14). In some fish species, populations of Gnrh neurons do not express Kiss receptors so cannot be directly regulated by Kiss as in mammalian models (9).

Observations such as these have accelerated research to uncover additional critical neuropeptides controlling reproduction (7, 15). For example, there are important stimulatory actions of neurokinin b (Nkb) on Gnrh neurons and directly on Lh and Fsh release in Nile tilapia (16). Yet, zebrafish with single and double mutations in the Nkb precursor genes *tac3a* and *tacb3b* exhibit completely normal fertility (17). Another peptide that has received attention is oxytocin (Oxt), also known as isotocin in teleosts. While implicated in Gnrh1 release and the Lh surge in rodents and humans (18, 19), *oxt* and *oxtr1* mutations in medaka disrupt female recognition of familiar males but ovulation, spawning success and fertility were not affected (20). Genetic manipulations of classical neuropeptides have largely yielded negative results in terms of impacts on fish reproduction. They are in marked contrast to the extensive pharmacological data demonstrating robust stimulatory effects of exogenously administered Gnrh, Kiss and Nkb on Lh release in several fish species (15, 21).

The possible exception is the neuropeptide secretoneurin (SN) that is derived from prohormone convertase-mediated processing of the large secretogranin-2 (Scg2) protein. Immunorectivity for SNa in nerve terminals in the anterior pituitary and in lactotrophs of goldfish supports the hypothesis that SN of both neuroendocrine and paracrine origin regulate pituitary Lh release (22, 23). Early studies in goldfish demonstrated that intraperitoneal (i.p.) injection of synthetic SN increases Lh release in vivo (24). In vitro studies established that SN could directly stimulate Lh production and release from dispersed goldfish pituitary cells in the absence of Gnrh (23, 25). Mouse LβT2 cells release both SN and LH in response to Gnrh1, and SN alone can stimulate Lh production and release, providing evidence for positive autocrine control (26). SN increases cyclic AMP and activates both protein kinase A and protein kinase C, causing the subsequent activation of extracellular signal-regulated kinase signaling pathways in LβT2 cells (26). These data suggest that SN action could be G-protein coupled, but the identity of the receptor remains elusive.

We recently reported that zebrafish carrying targeted frameshift mutations in the *scg2a* and *scg2b* genes have impaired sexual behaviors and reduced spawning success: only 1 in 10 double mutant couples could spawn (27). Significantly reduced expression of hypothalamic *gnrh3* and pituitary *lhb* subunit mRNA levels revealed the likely mechanisms underlying the disruption of spawning and failed ovulation. Intraperitoneal injection of the SNa but not SNb peptide into adults rapidly but only partially rescued reproductive defects (27). The subfertile *scg2a^-/-^; scg2b^-/-^* mutants resemble zebrafish and medaka mutant lines lacking Lh. In both species, mutation of the *lhb* gene caused female infertility linked to disrupted ovulation (28).

Most aspects of the evolution and physiology of secretograninergic neuroendocrine systems in normal animals of any species are still unknown. Therefore, we first conducted a broad phylogenetic analysis demonstrating that in the ancestor of teleost fishes, the duplication of the *scg2a* locus gave rise to the *scg2b* locus. We propose that SNa is the more ancient and conserved peptide. To establish the relationship of SNa and SNb to classical hormones we used our recently established sample extraction and sensitive mass spectrometry methods (29) to simultaneously measure periovulatory variations of 9 peptides and 5 steroids in the brain, pituitary and ovary. The strongest association we have uncovered was that brain SNa increased concomitantly with Gnrh3 prior to ovulation. Injection of synthetic SNa *in vivo* stimulates ovulation, placing SN amongst the other important neuroendocrine regulators.

## Results

### Phylogenetic analysis of the *scg2* family reveals gene duplication in teleosts

Most data available for SN are derived from studies in mammals (30,31), which have only one *scg2* gene, or from goldfish and zebrafish which have 2 genes, *scg2a* and *scg2b* (27,30). Yet only the SNa peptide, derived from prohormone convertase-mediated Scg2a precursor processing, has been studied in any detail. We set out to determine which gene is the more conserved of the two piscine forms. The *scg2* homologs from Chondrichthyes to Mammals including humans were identified (Fig S1). Likely a result of an incomplete genome assembly, we were only able to identify two truncated precursors in lampreys that had the SN domain. The data suggest that the Scg2 family may have originated in an ancestor of vertebrates. The *scg2* homologs found in numerous fish species included two copies in sturgeons, two in most teleost fishes and four in salmonids (Fig S1). Genome synteny analysis revealed the duplication signals in teleosts (Fig 1). For *scg2a*, the gene neighborhood is well conserved in all examined genomes and features two downstream genes, *wdfy1* and *gdn (serpine2)*. For the *scg2b* homologs, the gene neighborhood is marked by having two downstream genes: *cul3* and *mbnl1*. Between these two genomic loci, several genes are commonly shared and arranged, including the three central *scg2-ap1s3-dhrs12* genes. Several genes upstream of the *scg2a* and *scg2b* are also conserved, such as *rnf183, pax3, and epha4*, indicating that the duplication also involves the upstream region (Fig S2). Further gene insertion/deletion subsequently led to diversification at the *scg2b* genomic loci. In addition to the duplication event, we also found that the second duplication of salmon *scg2a* and *scg2b* involves 700 genes and 80 genes, respectively. For the sturgeons, we found the duplication involves a genome region containing about 40 genes. The data support that in the ancestor of teleost fishes, the duplication of the *scg2a* locus gave rise to the s*cg2b* locus, which produced two copies of the *scg2* gene. We propose that SNa is the more ancient and conserved peptide, which is equivalent to mammalian SN. This provides the foundation for our comparisons of a subset of classical reproductive hormones with the SN peptides.

**Fig 1.**
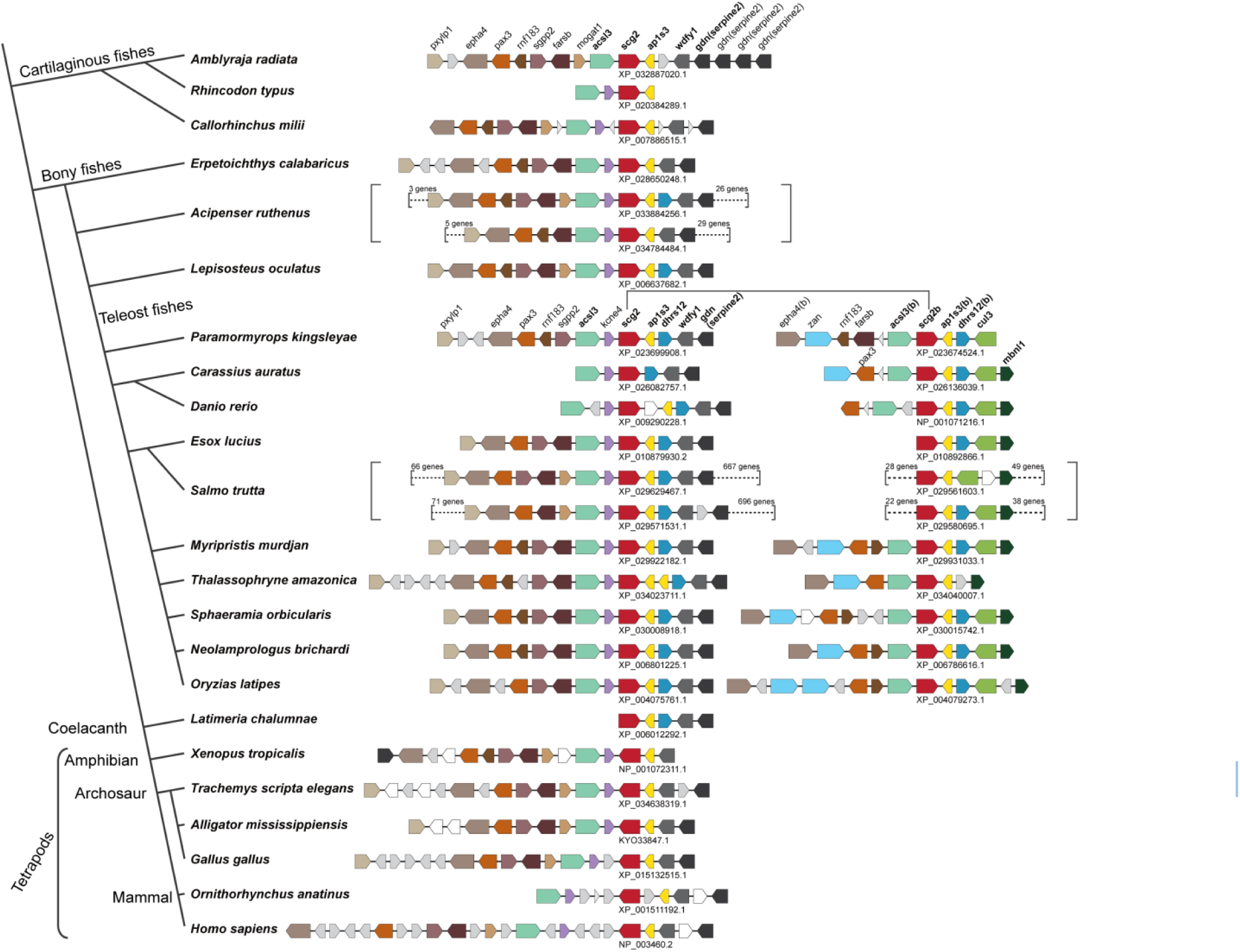
Genome synteny of *scg2* homologs from representative jawed vertebrates. The simplified tree topology, adapted based on Fig S1, is shown on the left side of the figure whereas their corresponding *scg2*-containing loci are shown on the right. Each orthologous gene group is represented by an arrowed box highlighted in different colors based on their annotations, shown above the locus. The genes encoding hypothetical proteins are shown in white boxes, non-coding RNAs in the gray boxes, and tRNA genes in triangles. The accession numbers of the *scg2* homologs are indicated below the loci. The genome duplication events found in *Acipenser ruthenus*, ancestor of teleost fishes, and *Salmo trutta,* determined based on Fig S2, are indicated by the red dots on the tree (left) and square brackets on the genome synteny (right). Only a part of duplicated regions in *Acipenser ruthenus* and *Salmo trutta* are shown.

### Neuroanatomical relationship between Scg2a/SNa and Oxt in brain and pituitary

We first determined the co-colocalization of Scg2a/SNa-ir (Fig 2A, D, G) and Oxt-ir (Fig 2B, E, H). This was prominent in preoptic magnocellular and gigantocellular neurons and fibers (Fig 2C, F). The double immunofluorescence was revealed in the cell bodies and in fine projections. Fibers displaying both Scg2a/SNa-ir and Oxt-ir can be seen entering the pituitary via the infundibular stalk (Fig 2G, H, I). Co-labelling was observed throughout the central portion of the pituitary and evident in Oxt-positive nerve terminals in the neurointermediate lobe (NIL) (Fig 2I). There are also SNa-positive/Oxt-negative terminals in the pars distalis of the anterior pituitary (Fig 2I). Strong Scg2a/SNa-ir was evident throughout the pituitary of Tg(*lhb*-RFP X *fshb*-eGFP) zebrafish (Fig 3A-C). Stained Scg2a/SNa fibers (Fig 3C, F) can also be seen in the proximal pars distalis (PPD) and rostral pars distalis (RPD). Gonadotrophs expressing Lh (Fig 3D, G) and Fsh (Fig 3E, H) were surrounded by Scg2a/SNa-ir. We found no evidence for colocalization of Scg2a/SNa in the gonadotrophs (Fig 3G, H, I).

**Fig 2.**
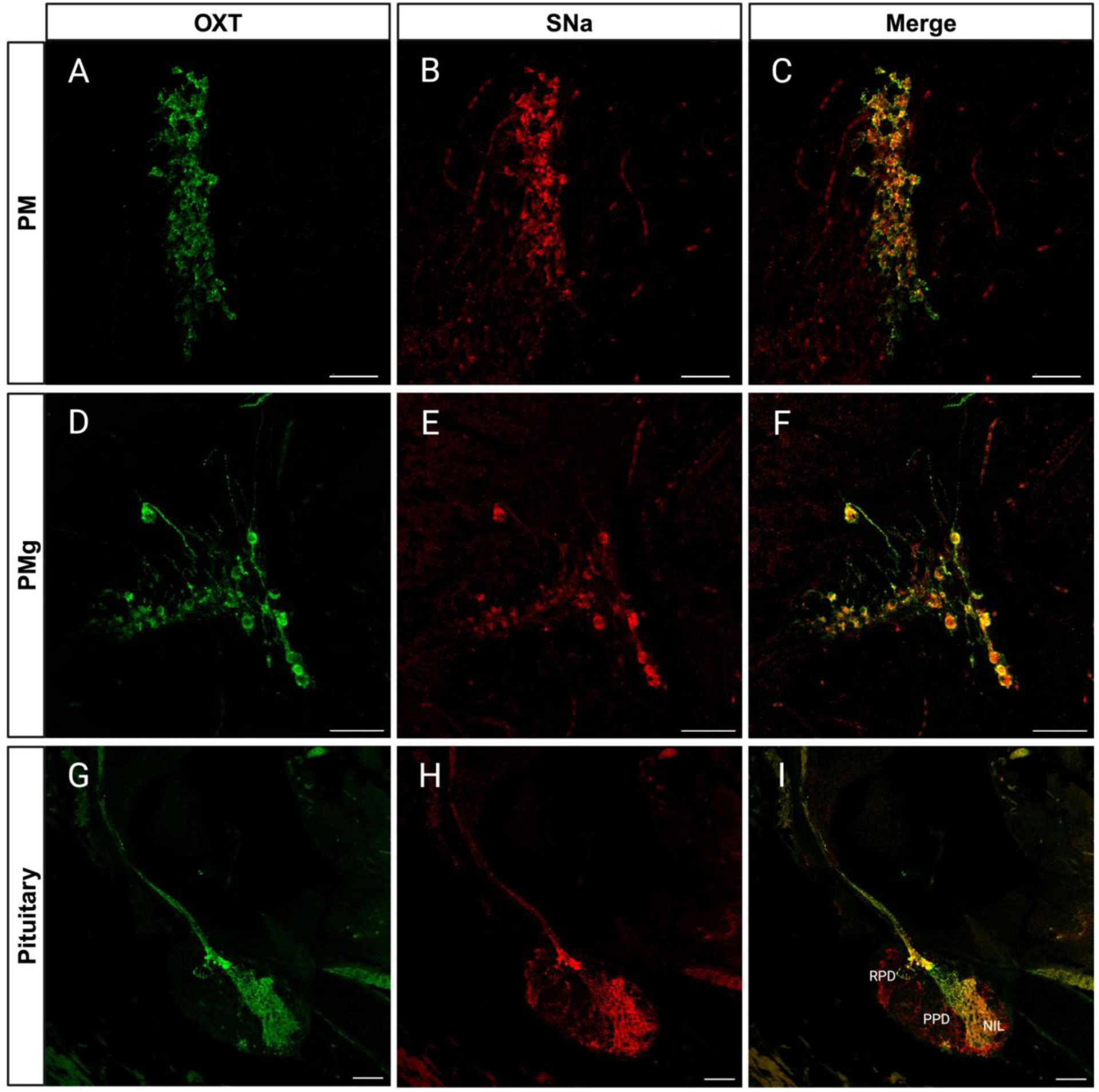
Immunocytochemical localization of Oxt-ir and Scg2a/SNa-ir in female zebrafish. Microscopy imaging of Oxt-ir (green) and Scg2a/SNa-ir (red) preoptic magnocellular (PM; panels A-C) and gigantocellular (PMg; panels D-F) regions, and pituitary (panels G-I). Merged images show colocalization (yellow-orange) of both immunoreactives in these paraffin sections. RPD, rostral pars distalis; PPD, proximal pars distalis; NIL, neurointermediate lobe. Panels A-F; 20X magnification, panels G-I; 10X magnification. Scale bars = 50 µm.

**Fig 3.**
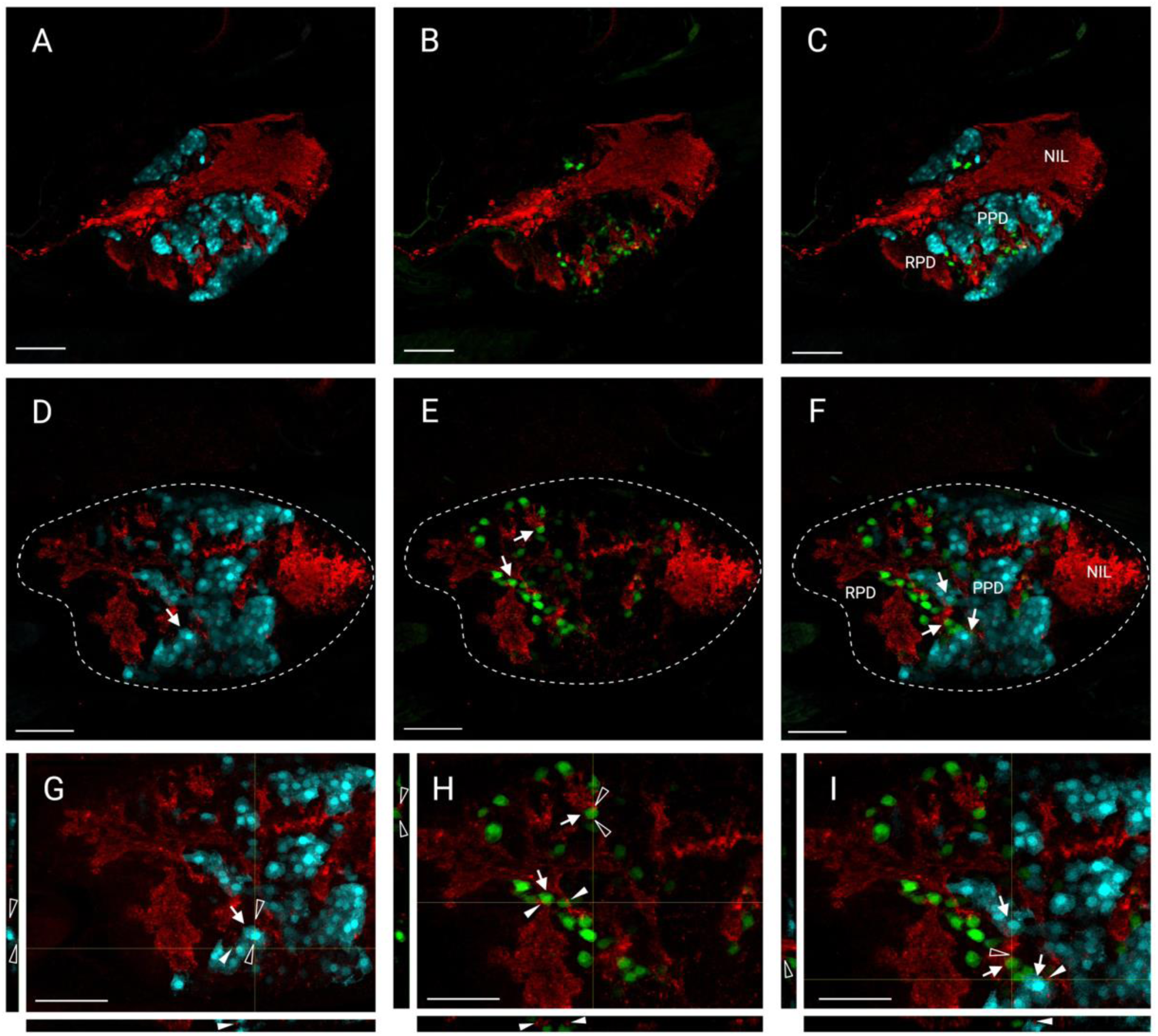
Distribution of Scg2a/SNa-ir in the pituitary of female Tg(*lhb*-RFP X *fshb*-eGFP) zebrafish. Confocal microscopy images obtained from cryosections (20 µm) show the distribution of Scg2a/SNa-ir (red), *lhb*-RFP (Cyan blue) and *fshb*-eGFP (green). The border of the entire pituitary gland is delineated with the white dashed lines, and solid white arrows (D,E,F) indicate cells of interest in magnified images (G,H,I). Optical slices in both planes are indicated by the faint yellow lines and can be seen to the left and bottom of the panels. Open arrowheads indicate SNa-ir around transgenic cells along the x-plane. Solid arrowheads indicate SNa-ir around transgenic cells along the y-plane. RPD, rostral pars distalis; PPD, proximal pars distalis; NIL, neurointermediate lobe. Scale bars = 50 µm.

### Periovulatory variations in Gnrh3 and Oxt

We sampled females at 02:00h (T1), 05:00h (T2) and 08:30h (T3). These were targeted because they are respectively the time points for: high pituitary *lhb* and *cga* mRNA levels (T1), the LH surge (T2) and ovulation (T3) (32–34). Reproductive neuropeptides including Gnrh2, Gnrh3, Kiss1, Kiss2, Oxt and Avp (arginine vasopressin, also known as vasotocin in fish) were detected in the samples of whole brain, pituitary, and ovaries from tissues harvested from individual fish (Fig 4 and Table S1). Levels of brain Gnrh3 and Oxt, and pituitary Oxt exhibited distinct patterns over the periovulatory period (Fig 4), while the other neuropeptides did not change (Table S1). Brain Gnrh3 increased 1.6-fold from T1 to T2 and then decreased significantly at T3 (p<0.05) (Fig 4A). Gnrh3 was detected in the pituitary, with a tendency for higher levels in some fish at T3 (Fig 4B). Moreover, Gnrh3 was detected in ovaries, although with no significant variation during the cycle (Fig 4C). As for Oxt, significant changes were observed in the brain and pituitary. Oxt variations in the brain (Fig 4D) and pituitary (Fig 4E) trended in opposite directions (Fig 4D). In the brain, Oxt was low at T1 and T2, but at T3, it increased significantly about 2.9 and 3.3-fold compared to the earlier sampling times (p<0.01). Pituitary Oxt decreased 74% from T1 to T3 (p<0.01) (Fig 4E). In ovaries, Oxt was below the detection limit (6.5 fmol/tissue), except for 1 of 10 samples (Fig 4F).

**Fig 4.**
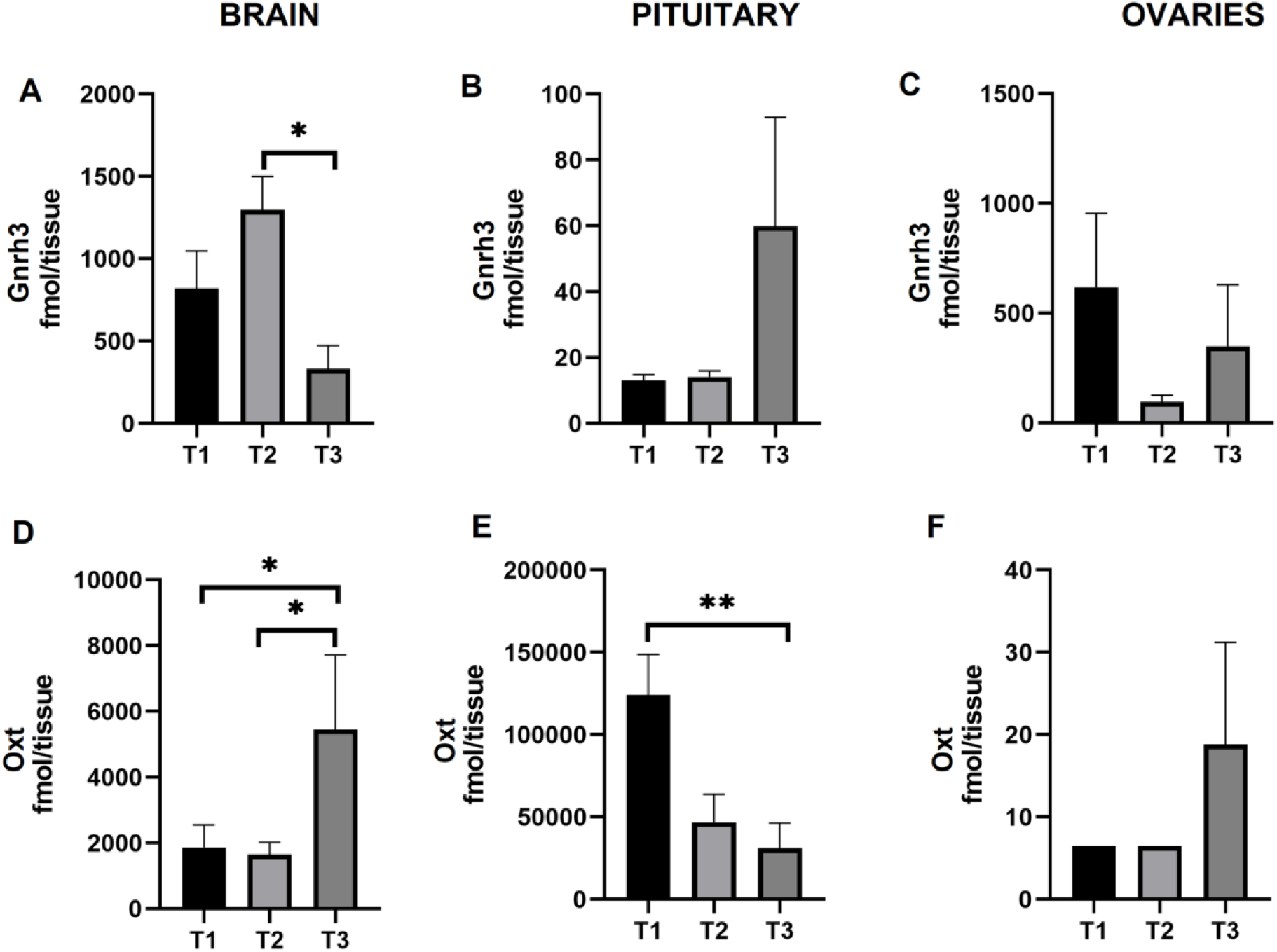
Gnrh3 and Oxt levels during the periovulatory period in brain, pituitary and ovaries of female zebrafish. The time points represent (T1) 02:00h, (T2) 05:00h and (T3) 08:30h. Gnrh3 in brain (A), pituitary (B), ovaries (C). Oxt in brain (D), pituitary (E), ovaries (F). Results are presented as mean+SEM (n=10). Note the different scales on the Y-axes. The Kruskal-Wallis one-way analysis of variance on ranks was performed followed by a Tukey test. *p<0.05. **p<0.01

### Periovulatory variations in estrone and estradiol levels

We simultaneously quantified estrone (E1), estradiol (E2), estriol (E3), testosterone (T), and 11-keto-testosterone (11-KT) with the neuropeptides (Table 1). In the brain, E1 levels decreased slightly from T1 to T2 (p>0.05) but increased 3.7-fold at T3 compared to T2 (p<0.01). In the pituitary, E1 levels peaked at T1 and decreased significantly from T2 (p<0.01). Significant variation in E2 was only evident in the brain. At T1 and T2, brain E2 was low but at T3 it increased 5.7- and 7.4-fold compared to T1 (p<0.05) and T2 (p<0.01), respectively. Levels of E3 decreased at T2 and T3 in all the tissues.

**Table 1.** Sex steroid levels (fmol/tissue) in the zebrafish periovulatory period. The time points represent (T1) 02:00h, (T2) 05:00h and (T3) 08:30h. ^a,b^Means ± SEM (n=10) with different superscripts are statistically different (p<0.05).

| Steroid |  | T1 | T2 | T3 |
| --- | --- | --- | --- | --- |
| E1 | Brain | 269.4±74.7 <sup>a, b</sup> | 168.0±48.5 <sup>b</sup> | 628.5±154.2 <sup>a</sup> |
|  | Pituitary | 594.7±117.5 <sup>a</sup> | 151.4±43.7 <sup>b</sup> | 146.8±53.4 <sup>b</sup> |
|  | Ovary | 643.7±337.1 | 363.1±192.6 | 446.6±232.9 |
| E2 | Brain | 137.0±30.6 <sup>a, b</sup> | 105.6±47.0 <sup>b</sup> | 781.1±194.6 <sup>a</sup> |
|  | Pituitary | 248.7±78.5 | 71.6±19.1 | 140.3±93.6 |
|  | Ovary | 313.8±258.7 | 22.6±11.6 | 57.1±36.0 |
| E3 | Brain | 1098.0±532.1 | 82.6±0.3 | 84.0±1.3 |
|  | Pituitary | 459.2±273.2 | 89.0±6.8 | 86.0±3.6 |
|  | Ovary | 596.5±344.6 | 246.0±163.4 | 82.9±0.9 |
| T | Brain | 11.9±3.1 <sup>a</sup> | 11.6±3.9 <sup>a</sup> | 1.1±0.0 <sup>b</sup> |
|  | Pituitary | 1.1±0.0 | 1.6±1.5 | 10.4±6.3 |
|  | Ovary | 4.3±2.1 | 1.1±0.0 | 1.1±0.0 |
| 11-KT | Brain | 50.3±5.7 <sup>a</sup> | 45.6±7.5 <sup>a, b</sup> | 21.5±8.8 <sup>b</sup> |
|  | Pituitary | 45.2±4.5 | 52.7±1.2 | 31.5±8.4 |
|  | Ovary | 73.2±42.4 | 62.6±14.9 | 35.2±7.4 |

### Brain androgen decreases while estrogens increase at ovulation

Periovulatory variations in T and 11-KT were only observed in the brain (Table 1). Brain T was relatively high at T1 and T2 but decreased 10.2-fold at T3 (p<0.05). Brain 11-KT levels at T3 were 43% of those measured at T1 (p<0.05). The ratio of E2 to T and that of E3 to E2 was also analyzed (Table 2). The brain E2/T ratio increased 33- and 10.9-fold at T3 compared to T1 and T3, respectivley. In contrast with the E2/T ratio, the E3/E2 ratio decreased significantly from T1 to T2 (p<0.05) and T3 (p<0.01). Ovarian T levels were generally low compared to 11-KT, and levels did not vary significantly during the periovulatory period (p<0.05). These data indicate that most of the ovarian T is converted to E2 and/or 11-KT.

**Table 2.**
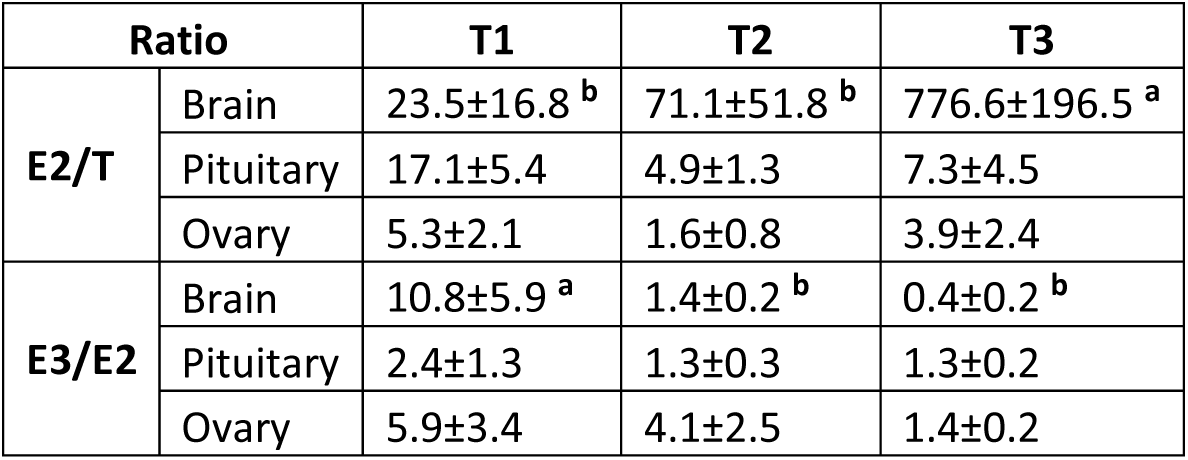
Ratios of E2/T and E3/E2 in the zebrafish periovulatory period. The time points represent (T1) 02:00h, (T2) 05:00h and (T3) 08:30h. ^a,b^Means ± SEM (N=10) with different superscripts are statistically different (p<0.05).

| Ratio |  | T1 | T2 | T3 |
| --- | --- | --- | --- | --- |
| E2/T | Brain | 23.5±16.8 <sup>b</sup> | 71.1±51.8 <sup>b</sup> | 776.6±196.5 <sup>a</sup> |
|  | Pituitary | 17.1±5.4 | 4.9±1.3 | 7.3±4.5 |
|  | Ovary | 5.3±2.1 | 1.6±0.8 | 3.9±2.4 |
| E3/E2 | Brain | 10.8±5.9 <sup>a</sup> | 1.4±0.2 <sup>b</sup> | 0.4±0.2 <sup>b</sup> |
|  | Pituitary | 2.4±1.3 | 1.3±0.3 | 1.3±0.2 |
|  | Ovary | 5.9±3.4 | 4.1±2.5 | 1.4±0.2 |

### Periovulatory variations in SNa1-34 and SNa1-14 support a potential role in ovulation

We employed untargeted proteomics and determined the amino acid sequences of SNa1-34 and SNb1-31 peptides. The existence of 5 fragmental peptides derived from SNa1-34 and 19 from SNb1-31 was established (Table S2). Of the smaller processed peptides, we selected SNa1-14 to investigate its function because its level is much higher than all other SNa or SNb-derived fragments. The HPLC-MS results revealed that the levels of the SNa1-34 and SNa1-14 peptides varied during the periovulatory period (Fig 5). In the brain (Fig 5A), SNa1-34 fluctuated significantly (p<0.01) and peaked at T2, which was 1.4- and 1.5-fold higher than at T1 (p<0.01) and T3 (p<0.01), resxpectively. Significant variations of SNa1-34 levels were also evident in the pituitary. SNa1-34 levels were similarly high at T1 and T2 (p>0.05) (Fig 5B) and declined by ∼50% at T3 (p<0.01). In the ovaries (Fig 5C), the SNa1-34 levels declined gradually from T1 to T3 (p<0.05). For SNa1-14, significant variations were evident in the brain and ovaries, but not in pituitary (Fig 5D-F). In contrast to SNa1-34, SNa1-14 levels increased gradually from T1 to T3 in both of brain and ovaries and peaked at T3 (Fig 5D and 5F). In the brain, SNa1-14 at T3 was 4.7-fold higher than that at T1 (p<0.05) while not statistically different than that at T2 (Fig 5D). In the ovaries, SNa1-14 levels were highest at T3 and respectively 2.7 and 3.1-fold higher than at T1 (p<0.01) and T2 (p<0.01) (Fig 5F). We also analyzed the ratio of SNa1-14 to SNa1-34 to investigate the major shift in SNa processing at ovulation. This was possible because we simultaneously measured all peptides from the tissues of individual animals. The SNa1-14/SNa1-34 ratios in the brain, pituitary and ovaries were all highest at T3 (Fig 5G-I). In brain, the SNa1-14/SNa1-34 was lowest at T2, rising significantly thereafter at T3 (p<0.01) (Fig 5G). The SNa1-14/SNa1-34 ratio in the pituitary was similar at T1 and T2 and increased significantly at T3 (p<0.01) (Fig 5H). In ovaries, the SNa1-14/SNa1-34 ratio increased gradually (Fig 5I), being different between T1 to T3 (p<0.01). The SNb peptide was detected in the brain, pituitary, and ovaries but there were no periovulatory changes (p>0.05) (Fig S3).

**Fig 5.**
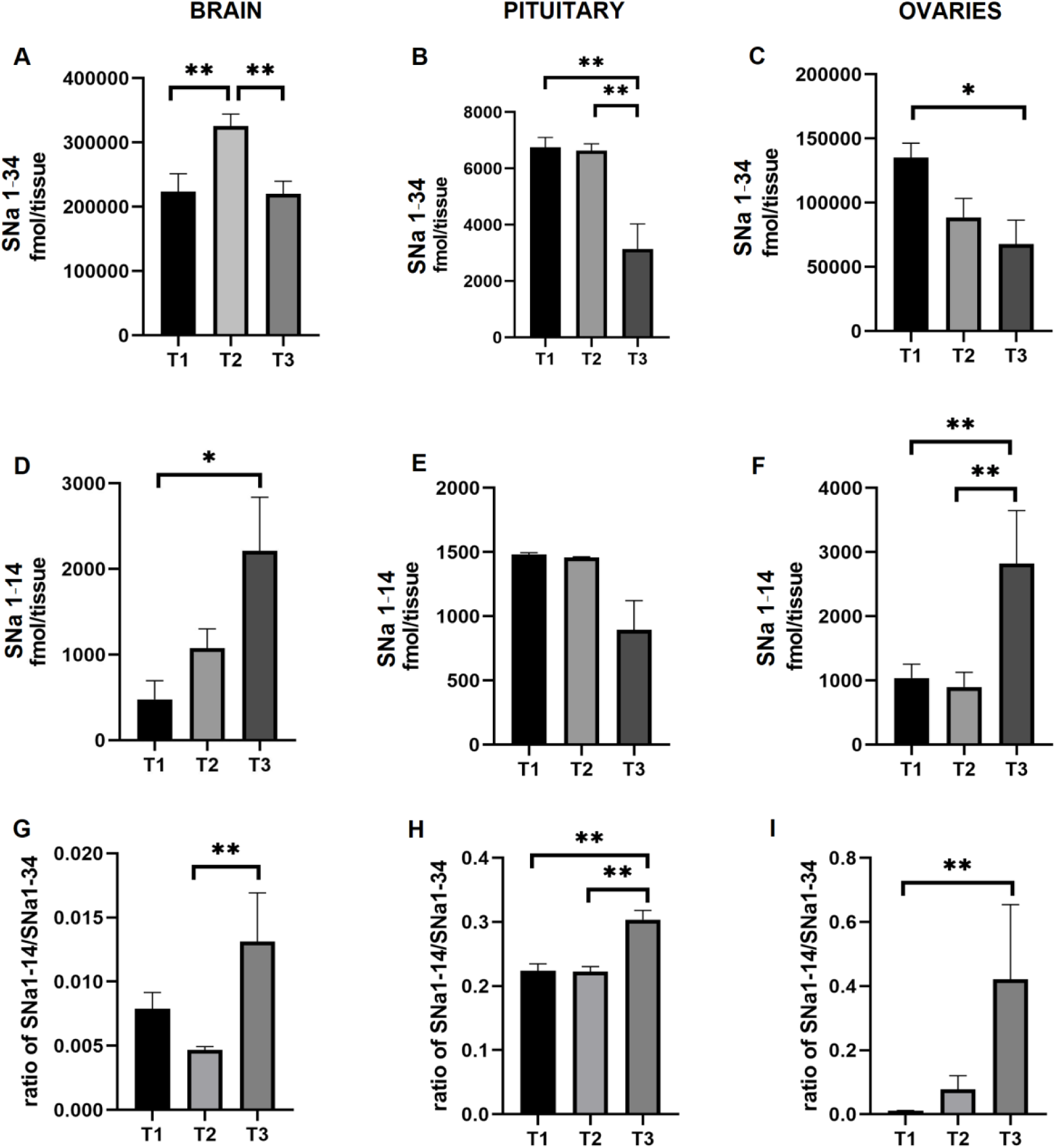
SNa levels vary during the periovulatory period in brain, pituitary and ovaries of female zebrafish. The time points represent (T1) 02:00h, (T2) 05:00h and (T3) 08:30h. SNa1-34 in brain (A), pituitary (B), and ovaries (C). SNa1-14 in brain (D), pituitary (E), and ovaries (F). Ratio of SNa1-14 to SNa1-34 in brain (G), pituitary (H), and ovaries (I). Results are presented as mean+SEM (n=10). Note the different scales on the Y-axes. The Kruskal-Wallis one-way analysis of variance on ranks was performed followed by a Tukey test; *p<0.05 **p<0.01.

### SNa levels in brain increase with Gnrh3 before ovulation

To investigate relations between SNa levels and the classical reproductive neuropeptides and sex steroids, we performed regression analysis (Table S3). Most R^2^ values between SNa and other hormones in the pituitary and ovaries were lower than 0.5 (Table S3). In marked contrast, SNa1-34 levels were strongly and positively related to levels of Gnrh3 in the brain (R^2^=0.7067). We also found that levels of SNa1-14 (R^2^=0.5765) were moderately related to Gnrh2 in the brain.

### SNa1-34 stimulates ovulation

The positive relation between SNa-related peptides and the Gnrh peptides suggested that either the phylogenetically conserved SNa1-34 and/or its SNa1-14 derivative peptide could be involved in the ovulatory cycle. Only SNa1-34 robustly induced ovulation in the afternoon outside the normal early morning ovulatory period, and in the absence of any males (Fig 6A). For females injected with SNa1-34, 11/18 (61%) ovulated by 6 h (p<0.01) post-injection, compared to 13/18 (72%) for those injected with the Lh analog hCG (p<0.01). In those females that ovulated, the % of ovulated eggs were similarly high in the SNa1-34 and hCG groups (p>0.05; Fig 6B). No fish ovulated following injections with saline or SNa1-14.

**Fig 6.**
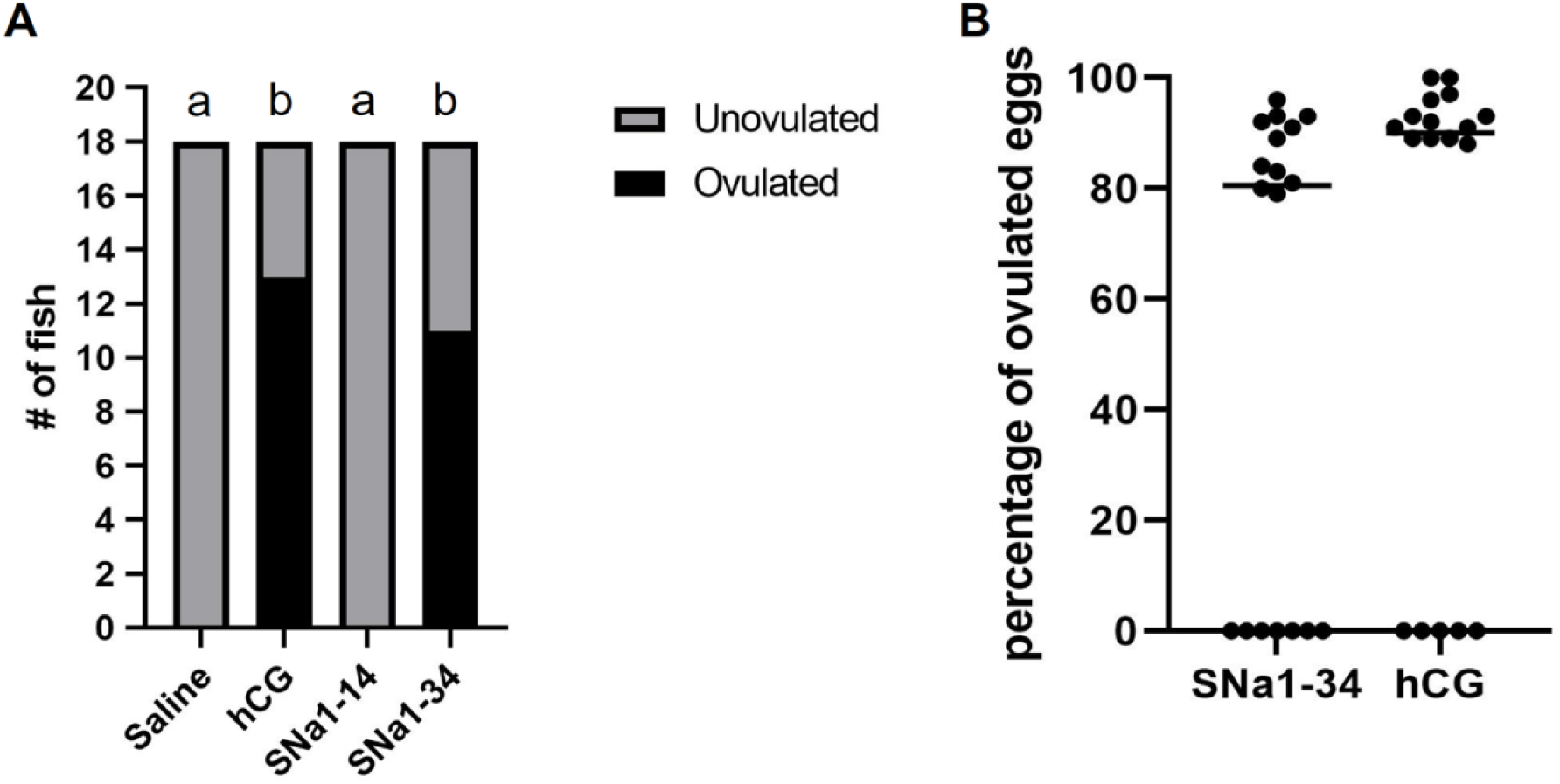
Ovulation of female zebrafish 6 h following i.p. injection with saline, SNa1-14, SNa1-34 and hCG. (A) The number of ovulating females was compared using Fisher’s exact test. Different letters (a,b) above the bars indicates a significant difference (p<0.01) in the numbers of ovulated females. (B) The percentage of ovulated eggs in female zebrafish injected with SNa1-34 or hCG. Individual levels are depicted along with median values (horizontal line; n=18); Mann-Whitney U-test; p>0.05.

### SNa1-14 and SNa1-34 differentially increase expression of key neuroendocrine genes in telencephalon and hypothalamus

We used droplet digital PCR (ddPCR) to quantify *gnrh2, gnrh3, oxt* and *avp* mRNA levels in telencephalon (Fig 7A-D) and hypothalamus (Fig 7E-H) harvested from individual females injected with either saline, SNa1-14 or SNa1-34. Except for *avp*, all the other genes in telencephalon were influenced significantly by treatment and time (Fig 7A-D). There was an overall effect of SNa1-14 and SNa1-34 injection to increase *gnrh2* (F=7.735, p=0.012), while time (F=3.515, p=0.103) and the treatment X time interaction were not significant (F=4.325, p=0.070) (Fig 7A). At 6 h post injection, telencephalon *gnrh2* in SNa1-14 injected females was 3.3-fold higher than those injected with saline (p<0.05). As for SNa1-34 treatment, the expression level of *gnrh2* increased 2.4-fold compared to saline injections and SNa1-34 treatment (p<0.05) (Fig 7A). For *gnrh3*, there were significant effects of treatment (F=8.390, p=0.021), but not time (F=3.723, p=0.095) (Fig 7B). Importantly, the treatment X time interaction (F=6.270, p=0.029), revealed that SNa1-34 had a time-dependent stimulatory effect on *gnrh3*. Levels of *gnrh3* in SNa1-34 injected females rapidly increased 8-fold at 3 h (p<0.05) but returned to control levels by 6 h post-injection. In contrast, SNa1-14 did not affect *gnrh3* in the telencephalon (Fig 7B). The SNa peptides exerted differential effects on *oxt* and *avp* in the telencephalon. Levels of *oxt* in telencephalon were affected by treatment (F=24.05, p<0.0001), time (F=6.340, p=0.039), and exhibited a significant treatment X time interaction (F=7.325, p=0.013) (Fig 7C). Injection of SNa1-14 and SNa1-34 resulted in a respective 6.9-fold (p<0.05) and 8.7-fold (p<0.05) increase in *oxt* at 3 h. The levels of *oxt* in fish injected with SNa1-34 at 6 h were not significantly different from saline injected fish (p>0.05) (Fig 7C) but were higher than SNa1-14 group (p<0.05). In contrast to *oxt*, the levels of *avp* were not affected by SNa1-14 or SNa 1-34 injection (F=4.250, p=0.076) at any time (F=2.601, p=0.151) nor were there any treatment X time interactions (F=2.385, p=0.163) (Fig 7D).

**Fig 7.**
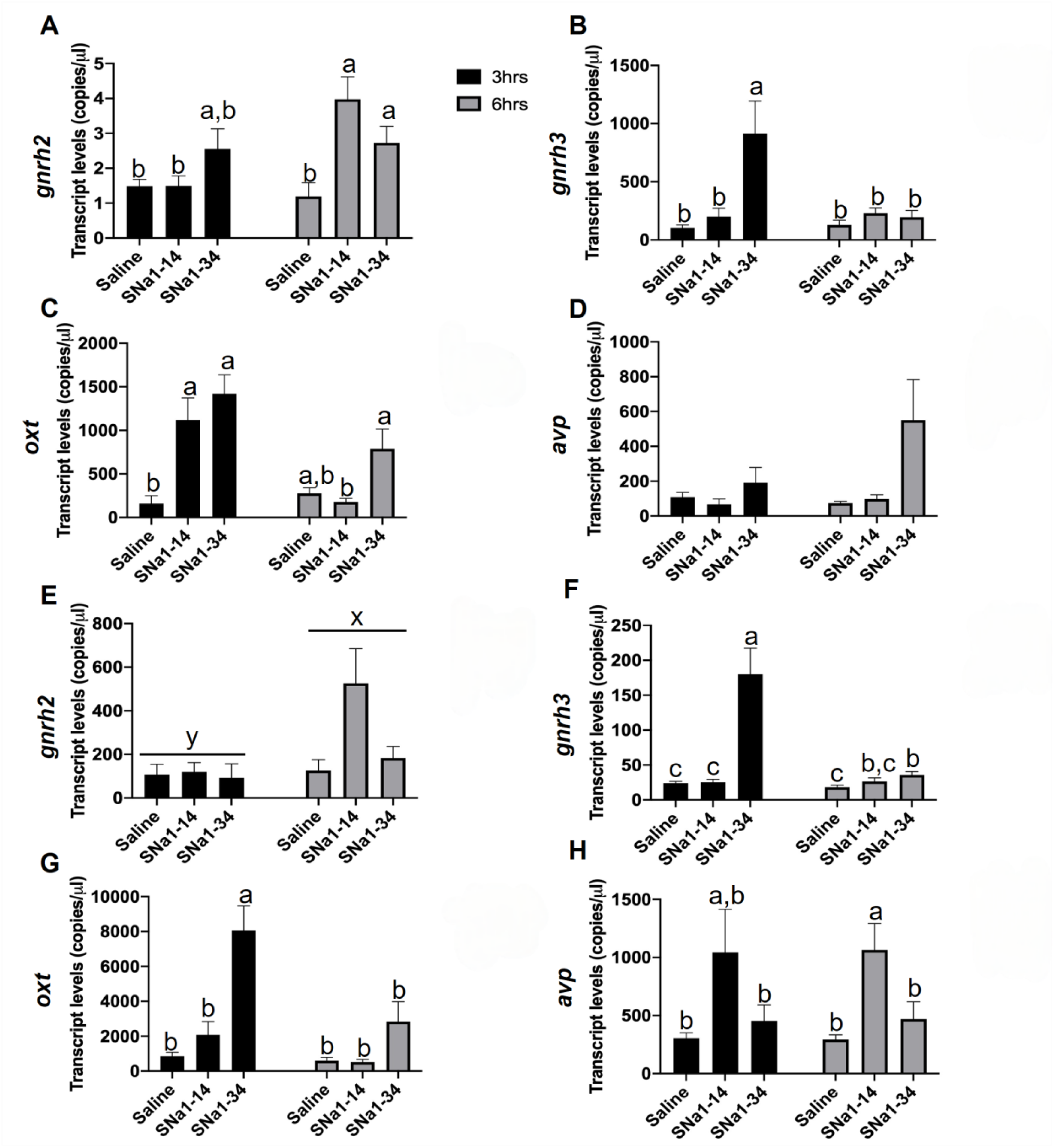
Effects of SNa1-14 and SNa1-34 on reproductive neuropeptide in telencephalon (A-D) and hypothalamus (E-H) in female zebrafish. Telencephalon expression of *gnrh2* (A)*, gnrh3* (B), *oxt* (C), *avp* (D) and hypothalamus expression of *gnrh2* (E)*, gnrh3* (F), *oxt* (G), *avp* (H) are presented (mean+SEM; n=8). In panel E, the different superscripts (x,y) indicate a significant main effect of time (F=10.14, p=0.015). In other panels, means with different letters (a,b) indicate a significant difference (Two-way ANOVA; p<0.05).

The same genes were quantified in the hypothalamus (Fig 7E-H), but the expression patterns were different than those observed in the telencephalon. Levels of *gnrh2* were not affected by treatment (F=4.248, p=0.072) and there was no treatment X time interaction (F=3.381, p=0.073). There was an overall trend for *gnrh2* to be higher at 6 h than 3 h (F=10.14, p=0.015) (Fig 7E). Injection of SNa1-34 but not SNa1-14 increased hypothalamic *gnrh3* (F= 19.98, p=0.002) in a time-dependent manner (F= 12.39, p=0.009), with a clear treatment X time interaction (F= 13.40, p=0.007) (Fig 7F). At 3 h post SNa1-34 injection, *gnrh3* increased 7.3-fold. By 6 h, *gnrh3* levels decreased significantly (p<0.05) although it remained somewhat higher (p<0.05) in SNa1-34 injected fish compared to those injected with saline (Fig 7F). In a pattern like that observed for *gnrh3*, injection of SNa1-34 but not SNa1-14 increased hypothalamic *oxt* (F= 28.39, p<0.0001) in a time-dependent manner (F= 8.489, p=0.022), with a treatment X time interaction (F= 5.800, p=0.033) (Fig 7G). At 3 h post-injection of SNa1-34, the level of *oxt* increased 8.2-fold (p<0.05), then declined significantly (p<0.05) to basal levels at 6 h. Injection of SNa1-14 but not SNa1-34 had an overall stimulatory effect on hypothalamic *avp* (F= 6.116, p= 0.040) to a similar level at both times (F= 0.1825, p=0.682) (Fig 7H). No treatment X time interaction (F= 0.1795, p=0.703) was evident for *avp*. At 3 and 6 h following SNa1-14, *avp* was increased 3.4-fold (p>0.05) and 3.5-fold (p<0.05), respectively, compared to the saline groups.

### SNa1-34 but not SNa1-14 increases the expression of pituitary Lh subunits and key ovarian genes

Transcript levels of *cga, lhb, lhcgr* and *npr* were increased following SNa1-34 injection. There was a significant main effect of treatment to increase the levels of pituitary *cga* (F=15.23, p=0.002) that was independent of time (F=0.1142, p=0.745) and there was no treatment x time interaction (F=1.530, p=0.256) (Fig 8A). Injection of SNa1-34 increased *cga* 9-fold and 3-fold at 3 h and 6 h, respectively, compared to saline controls. In contrast, SNa1-14 did not influence *cga*. Levels of *lhb* in pituitary were significantly increased following injection of SNa1-34 (F=8.898, p=0.019) (Fig 8B) but it was not affected by time (F=0.4146, p=0.540) and no treatment x time interaction was evident (F=0.6299, p=0.460). Transcript levels of *lhb* were 6.6-fold and 3-fold higher at 3 h and 6 h, respectively, compared to saline injected fish. SNa1-14 did not influence *lhb*. Transcript levels of pituitary *fshb* were not significantly different between SN or saline injected fish (Fig 8C). The stimulatory effects on *cga* and *lhb* at the level of the pituitary, lead us to examine gene markers of ovarian activation. Injection of SNa1-34 increased ovarian levels of *lhcgr* (F=4.392, p=0.036), a time-dependent effect was observed (F=6.113, p=0.042), yet no treatment X time interaction was evident (F=2.551, p=0.139). Levels of *lhcgr* were 2.5-fold higher in the SNa1-34 group at 3 h compared to the saline group (p<0.05), then they returned to basal levels after 6 h. In contrast, no differences in the levels of *lhcgr* were observed between SNa1-14 or saline injected fish (Fig 8D). Levels of ovarian *cyp19a1a* remained stable across treatments (F=0.603, p=0.504), time (F=3.448, p=0.105) and treatment X time interaction (F=2.892, p=0.098) (Fig 8E). Injection of SNa1-34 increased ovarian *npr* (F=1.733, p=0.036) (Fig 8F). There was a 3.3-fold increase in ovarian *npr* at 3 h (p<0.05), there was no main effect of time (F=2.319, p=0.171) and the treatment x time interaction (F=1.279, p=0.304) was not evident. Levels of *npr* returned to basal levels after 6 h.

**Fig 8.**
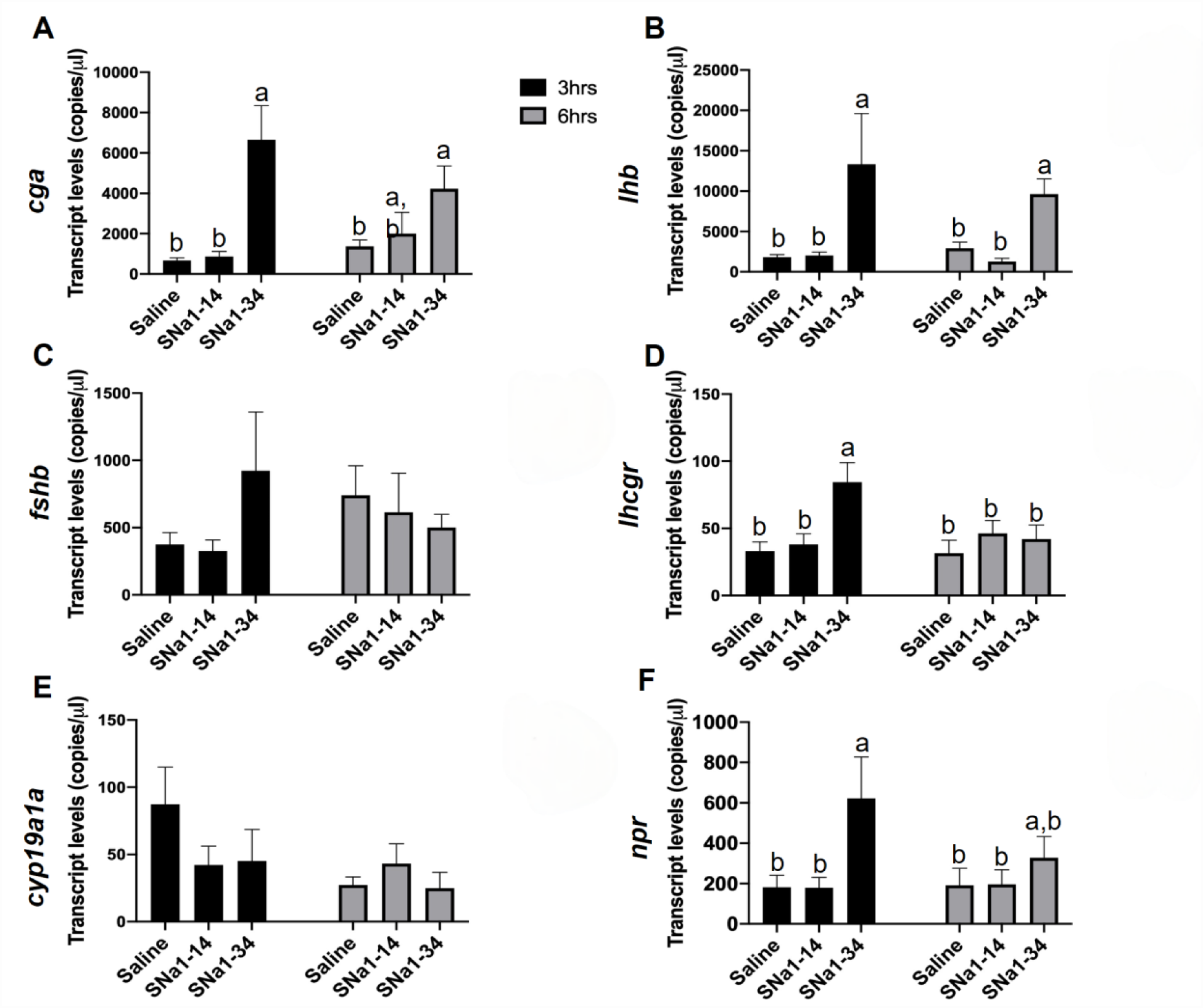
Effects of SNa1-14 and SNa1-34 injection on reproduction-related gene expressions in pituitary (A-C) and ovary (D-F) in female zebrafish. Transcript levels of *cga* (A)*, lhb* (B), and *fshb* (C) in pituitary and levels of *lhcgr* (D)*, cyp19a1a* (E), and *npr* (F) in ovary are presented (mean+SEM; n=8). Means with different letters (a, b) indicate a significant difference (Two-way ANOVA; p<0.05).

## Discussion

We provide evidence for periovulatory variations in peptides and steroids in tissues of female zebrafish. These data are significant for three main reasons: 1) we report on the specific relationship between preovulatory increases in brain SNa1-34 and Gnrh3 levels; 2) a single i.p. injection of SNa1-34 robustly stimulates key genes at each level of the HPG axis; and 3) treatment with SNa1-34 leads to ovulation. Figure S5 summarizes key findings and presents a working model for the role of SNa in zebrafish reproduction.

There are about 20 stimulatory neuropeptides along with classical neurotransmitters likely involved in the neuroendocrine control of teleost reproduction (15, 35), but very few of these neurohormones have ever been measured in naturally ovulating fish of any species. The main motivation to carefully document changes in the classical HPG hormones was to compare them to the Scg2-derived peptide SN. Genome synteny analysis indicated that the duplication of the *scg2a* locus gave rise to the s*cg2b* locus, which produced two copies of the *scg2* gene. We conclude that SNa in teleosts is the more ancient and conserved peptide, which is equivalent to mammalian SN.

To establish a neuroanatomical context, immunocytochemical studies revealed neuronal fibers staining for SNa projected from the POA, passed through the pituitary stalk, and terminated largely in the NIL. Immunoreactive SNa colocalized with Oxt in cell bodies in the preoptic area and fibers in the NIL. This relationship has also been observed in rats (36) and goldfish (22) implying it is phylogenetically conserved. In female medaka, a subset of preoptic neurons express neuropeptide B. Thus, named as female-specific, sex steroid-responsive peptidergic (FeSP) neurons, it has recently been shown that they also express *scg2a* (37). In situ hybridization for *scg2a* with *oxt* or *avp* showed that nonapeptide-expressing neurons in the female medaka preoptic area also co-express *scg2a*. The immunocytochemical colocalization of SNa with OXT implicates SN in the control of reproduction (18, 19) and in sociosexual behaviors (38,39). Some SNa-positive/Oxt-negative fibers were evident in the PPD of the zebrafish pituitary. We observed this SNa-ir near gonadotrophs but did not reveal any evidence for localization in them in the pituitaries of the Tg(*lhb*-RFP x *fshb*-eGFP) line. Immunoreactivity for SN-ir appears as small boutons near the gonadotrophs and suggests that some Lh cells receive innervation by neurons producing Scg2a/SNa.

Next, we used our sensitive HPLC-MS methods (29) for comparisons of classical reproductive hormones with the SN peptides. We simultaneously quantified 9 neuropeptides and 5 steroids in brain, pituitary, and ovary samples from individual female zebrafish. The results revealed that Gnrh3 and Oxt, and E1, E2, T and 11-KT varied during the periovulatory period. In contrast, Gnrh2, Avp, Kiss1 and Kiss2 and E3 were relatively stable. SNa-related peptides also exhibited periovulatory changes, but SNb did not.

GnRH is an important neuropeptide regulating ovulation by stimulating the Lh surge. Different from mammals, zebrafish produces both Gnrh2 and Gnrh3. Extensive neuroanatomical, physiological and pharmacological data for teleosts support the importance of the multiple Gnrh forms for Lh synthesis, release, and ovulation (7, 8,9,35,40). Detection of Gnrh2 and Gnrh 3 in the brain and pituitary indicates that both are potentially hypophysiotropic in zebrafish. The variations we report for Gnrh3 support its role in ovulation. The Gnrh3 levels in brain were highest at T2, around the time of the Lh surge, preceding ovulation at T3. While, this may have been predictable, Gnrh3 and the other peptides have never been specifically quantified by MS previously. In contrast to brain, Gnrh3 levels in the pituitary and ovary did not change significantly over the periovulatory period.

Pituitary Oxt was highest at T1 when the pituitary *lhb* mRNA levels are reportedly high (32), whereas Oxt in brain increased significantly at T3, the time of ovulation. Although much evidence is lacking, our results indirectly support a role for Oxt in the periovulatory period. While Oxt has a minor stimulatory effect on Lh release from dispersed goldfish pituitary cells in vitro (41), it more robustly enhances Lh in the ricefield eel (42). On the other hand, normal *oxt* gene function is required for mate choice based on familiarity recognition in medaka (20). Mutation of *oxt* or *oxtr1* did not affect spawning success in these medaka, supporting the concept that Oxt modulates sociosexual behaviors rather than ovulation itself (20).

Ovarian E2 and E3 levels were highest at T1, but decreasing thereafter, being lowest at T3. At T1 there was high variation between females, so there were no statistically significant variations in ovarian estrogens evident. Intraovarian E2 (and presumably also E1) plays a key inhibitory role via G protein-coupled estrogen receptor controlling the onset of oocyte maturation and meiotic cell-cycle progression (43). After the ovulatory Lh surge, ovarian steroidogenesis favors the increased synthesis of a maturation-inducing steroid (MIS, e.g., 17α,20β-dihydroxy-4-pregnen-3-one and related progestins) that induces final maturation of the oocyte via nongenomic actions involving membrane progesterone receptors (44). Ovarian T levels were generally low, and while 11-KT was higher, levels did not vary significantly over the study. This implies that most of the T is converted to E2 and 11-KT.

The surprising results of steroid analysis relate to brain and pituitary. The highest levels of pituitary E1 were found at T1, 6.5 hrs before ovulation. The highest levels of brain E1 and E2 were noted for T3 at ovulation, and coincident with the lowest brain T levels. Calculation of the E2/T ratios for brain indicate a major shift to E2 production at ovulation. That the brain E2/E3 ratios were their lowest at ovulation suggests a decrease in E2 metabolism, contributing to high brain E2 at T3. Brain tissue steroid levels reflect both uptake from the periphery, and the estrogen synthesis capacity of teleost radial glial cells that exclusively express cyp19a1b/aromatase B (45). Recently, we reported that brain E2 levels in *cyp19a1b ^-/-^* mutants are 50% of levels in wildtype females (46,47), indicating approximately equal contributions of E2 from central and peripheral sources. Radial glial cells synthesizing estrogens surround the preoptic Oxt and Avp neurons that co-express SNa (22,46,47). In female medaka, estrogens regulate *scg2a* in preoptic FeSP neurons (37). Estrogens can also increase neuronal firing rates in reproductively relevant brain circuits in midshipman fish within minutes (48), or in Gnrh1 neurons of medaka within 2 days (49). Thus, the observed rapid shift in steroid dynamics in the female zebrafish brain is on a physiologically relevant timescale. Levels of E1 and E2 were significantly increased in the brain at T3. It is possible that this increased brain estrogen level could regulate post-ovulatory spawning activity. We have shown that *cyp19a1b^-/-^* mutant females have fish have reduced hypothalamic *oxt* and *avp* mRNA levels, delayed oviposition and reduced offspring survival (46,47).

Peptidomic analysis revealed the production of two major SNa-related peptides (SNa1-34 and SNa1-14) and SNb1-31, plus other quantitatively minor fragments. In comparing zebrafish SNa with bovine SN processing (50), we found a similar cleavage pattern. We hypothesized that some SN peptides could be biologically active after cleavage. To confirm the existence of the candidate fragments, we synthesized and purified SNa1-34, SNa1-14 and SNb1-31 and generated standard compounds for HPLC-MS. Differing periovulatory variations were observed for SNa1-34 and SNa1-14, suggesting that they play different roles. SNa1-34 was highest at T1 and T2 when the respective expected peaks in *lhb* mRNA and the Lh surge occur. A single SNa1-34 injection in subfertile *scg2a-/-; scg2b-/*- zebrafish enhances spawning by ∼3 fold, like hCG injection in the same mutant line (27).

Levels of SNa1-14 peaked at ovulation in both the brain and ovaries. Since there is no related research on SN fragments it is tempting to comment on the importance of these observations. The SNa1-14 to SNa1-34 ratio significantly increased in all three tissues at ovulation, indicating enzymatic processing at this time. Ovary levels of SNa1-34 decreased at ovulation, while SNa1-14 increased, further indicating a shift in SNa1-34 processing. Levels of *scg2a* have been quantified by PCR in goldfish ovaries (51) and *scg2a* and *scg2b* detected by RNASeq in zebrafish ovarian follicles (52). The ovary is thus a likely additional production site for Scg2 and/or its derived peptides. The role of ovarian SN is a matter of speculation, but it is now among the many other reproductive neuropeptides produced there. These include kisspeptin, Gnrh and nonapeptides we report here, some of which have demonstrated paracrine roles in gonadal function (53,54). In humans, nonhuman primates and rodents, hCG induces *scg2* mRNA and protein in granulosa cells, and SN added to human ovarian microvascular endothelial cell cultures induced angiogenesis, so SN has a role in the ovulatory process (55). Much remains to be discovered concerning the function of locally produced gonadal Scg2/SN.

Noteworthy is the observation that brain SNa and Gnrh3 both peaked at T2. This precedes ovulation by ∼3.5 h, implying a role for SNa1-34 in inducing the Lh surge. This is supported by the observation that i.p. injection of SNa1-34 increases *gnrh3, lhb,* and *cga* within 3 h (27). Note also that SNa can induce *lhb* mRNA and/or Lh release in 3-6 h from dispersed pituitary cells in goldfish and grouper (23, 56). In single identified goldfish gonadotrophs in culture, SNa application provokes a rapid increase in intracellular calcium, demonstrating direct cellular actions (23). In goldfish, SNa injection *in vivo* increased serum Lh in 30 min in animals pretreated with a dopamine antagonist (24). Thus, SN actions in the pituitary could also contribute to the Lh surge. Regression analysis indicated that Gnrh2 changes were only weakly related with SNa1-34, and moderately associated with SNa1-14. Yet, the lack of variation of Gnrh2 in the periovulatory period suggests that Gnrh2 is not directly involved in ovulation, at least during the period we assessed. Gnrh2 mutant zebrafish spawn normally but oocyte quality is reduced (57). In marked contrast with SNa, the SNb-related peptide levels did not vary in the periovulatory period in any tissue. This was surprising, since *scg2b* mutant zebrafish have decreased spawning rates compared to wildtype; however, SNb injection does not rescue spawning in these subfertile mutants (27). The role of the numerous other Scg2b-derived peptides in reproductive processes remains to be investigated.

We suspected that one of the major SNa forms may stimulate ovulation, so this was assessed in fish i.p. injected with synthetic SNa1-34 or SNa1-14 and compared to those injected with hCG. Human chorionic gonadotropin is a Lh analog that activates the key Lh-dependent pathways in ovaries (58) and stimulates ovulation in normal and *scg2a-/-; scg2b-/*- double mutant zebrafish (27). So hCG injection serves to mimic the Lh surge. A single injection of hCG induced ovulation in the female zebrafish. Injection of SNa1-34 induced a robust ovulatory response indistinguishable from hCG. In contrast, ovulation was not induced by the SNa1-14 fragment that is generated from natural processing of SNa1-34.

To assess the underlying neuroendocrine mechanisms for SNa-induced ovulation, we quantified the transcript levels of key genes in the HPG axis known to positively regulate reproduction. SNa1-34 robustly increased *gnrh3* in telencephalon and hypothalamus after 3 h, returning to baseline by 6 h. No effects of SNa1-14 on *gnrh3* transcript levels were evident. We found that SNa1-14 and SNa1-34 injection increased *gnrh2* transcript levels in the telencephalon at 6 h. Both Gnrh3 and Gnrh2 stimulate Lh release in fish (7, 8,15). Analogs of Gnrh3 in particular, have been used extensively for ovulation induction in numerous teleost species (59). Therefore, it is plausible that SNa1-34 participates in the Lh surge by stimulating Gnrh3 neurons, already shown through laser ablation studies to be critical for the ovulatory process in zebrafish (60). In addition to Gnrh, a lesser-known contributor to the Lh surge is Oxt (18,19). Ox**Error! Bookmark not defined.**t also stimulates Lh release in goldfish (41). In our female zebrafish, levels of *oxt* mRNA in telencephalon were increased following SNa 1-14, and SNa1-34 injection after 3 h but only SNa1-34 increased *oxt* levels at 3 h in the hypothalamus. Levels of Oxt peptide were highest in the pituitary before the time of the Lh surge, and highest in the brain at the time of ovulation, so it is possible that SNa1-34 through effects on Oxt could modulate reproductive outcome.

These stimulatory effects of SNa1-34 on *gnrh3* and *oxt* in brain tissues led us to also investigate effects on gonadotropin subunit gene expression in the pituitary. The most striking effect of SNa1-34 was the robust and concurrent 10- and 7-fold increase in *cga* and *lhb,* respectively, at 3h. The *cga* and *lhb* levels were also elevated at 6 h but not as high as at 3 h post-injection. No significant differences were observed in any of the gonadotropin subunits following injection of SNa1-14 or for the levels of *fshb* in fish treated with SNa1-34. The Lh surge drives ovulation and is indispensable in zebrafish reproduction. Female zebrafish with mutations of *lhb* are infertile because of the failure to ovulate (28). Female zebrafish carrying frameshift mutations in the *scg2a* precursor (27) have a very similar phenotype to the *lhb* mutants (28). Both have well-formed ovaries, but do not ovulate. Moreover, pituitary *cga* and *lhb* levels were decreased by >90% in *scg2a* mutant females (27). A range of physiological, pharmacological, and gene mutational studies in fish indicate that Fsh is important for oocyte development from primary ovarian growth to cortical alveolar stages, whereas Lh drives maturation and ovulation (58). Therefore, SNa1-34 injection robustly activates hypophysiotropic neurons and Lh-producing gonadotrophs, consistent with effects on ovulation.

To determine if brain and pituitary gene expression changes induced by SNa1-34 injection were associated with altered ovarian function, we measured the expression of ovarian *lhcgr* and *npr*. The main effects of SNa1-34 were increased levels of *lhcgr* and *npr* at 3 h post injection, coinciding with the increases in pituitary *cga* and *lhb*. Co-activation of Lh-producing gonadotrophs and ovarian Lh receptors is the hallmark of the ovulatory process (61,62). Moreover, ovarian *npr* was also increased by SNa1-34 at 3 h post-injection. Nuclear progesterone receptor is important for ovulation (58,63). Both Lhcgr and Npr are critical to the transformation of oocytes from the maturational to ovulation stages (58). In zebrafish, *npr* is stimulated by Lh, and *npr* is highest in full-grown stage ovarian follicles. Female zebrafish with mutated *npr* are infertile due to ovulation defects (33). Thus, increased *gnrh3* in brain, *lhb* in pituitary and *npr* in ovary following SNa1-34 injection are strong indicators of ovulation induction.

We have provided evidence that SNa1-34 plays a critical role in female reproductive processes. Natural variations in SNa1-34 associated with changes in well-known stimulatory neuropeptides in the brain have been quantified for the first time in any species. Increased SNa1-34 coincided with increased Gnrh3 at the time of the Lh surge, preceding ovulation. We demonstrate that injection of SNa1-34 induces ovulation, as does the Lh analog hCG. This effect of SNa1-34 provoked timely increases in key genes at the level of brain, pituitary, and ovary, commensurate with the ovulatory response. Together these data in normal animals along with those obtained in anovulatory female zebrafish harboring frameshift mutations in *scg2* genes (27), places SN amongst other important neuroendocrine regulators.

## Material and Methods

### Genome synteny analysis

Details on methods and results of sequence searches and phylogenetics analysis are in the SI. We extracted the upstream and downstream gene neighbors of the *scg2* homologs from the NCBI genome database. Protein sequences of the gene neighbors were clustered based on the sequence similarity through the BLASTCLUST program (ftp://ftp.ncbi.nih.gov/blast/documents/blastclust.html) and assigned with a common gene name which was extracted from the NCBI database. The gene clusters that are present in most of the vertebrate species were further selected for phylogenetic analysis.

### Experimental animals and husbandry

Wildtype zebrafish (AB strain) were bred and kept in the University of Ottawa Aquatics core facility at 28°C on a 14:10h light/dark cycle. All the fish were fed twice a day. The *lhb*-RFP and *fshb*-eGFP transgenic zebrafish described in detail previously (64) were crossed to obtain the Tg(*lhb*-RFP X *fshb*-eGFP) line. Sexually mature zebrafish at the age of 6-9 months post-fertilization (mpf) were selected and separated by sex for least two weeks to synchronize the reproductive cycles before the experiment. All animal experiments followed the guidelines of the Canadian Council on Animal Care for the use of animals in research, approved by the University of Ottawa Animal Care Protocol Review Committee. We have adopted the zebrafish gene/protein nomenclature conventions according to https://zfin.atlassian.net/wiki/spaces/general/pages/1818394635/ ZFIN+Zebrafish+Nomenclature+Conventions. Of note, the recently determined zebrafish nomenclature for *avp*/Avp and *oxt*/Oxt contrasts the historically used nomenclature of *avt*/Avt and *ist*/Ist. This change highlights the homologous nature of the nonapeptide genes and their protein.

### Immunofluorescence microscopy

After anesthesia and sacrifice of zebrafish in an ice bath, the heads were removed and incubated into 4% paraformaldehyde overnight at room temperature. Samples were washed in PBS (2 X 20 min) for twice and decalcified in 0.5M EDTA at room temperature for 5-7 days. Details of paraffin and cryosectioning are presented in the SI. The specificity of the guinea pig-anti-OXT antiserum (22, 65) in fish has been previously reported. The rabbit-anti-SNa antiserum was generated and extensively validated in our lab (22,25,66). It has been shown using western blotting that it will recognize the Scg2a precursor protein and any processed fragment containing the SNa epitope (66). Therefore, we refer to this as Scg2a/SNa immunoreactivity. The epitope (YTPQKLATLQSVFEE) recognized by the SNa antibody is identical to the central 15 amino acids in zebrafish SNa1-34 (Table S2) and does not cross-react with SNb1-31 (67). Following these validated procedures, sections were incubated with guinea pig-anti-OXT antiserum (Abcam, cat#ab228508) and rabbit-anti-goldfish SNa (1:1000) overnight at 4°C. The next day, sections were washed in PBS (2 x 20 min), then incubated at RT for 2 h with goat anti-Rabbit IgG (H+L) Alexa Fluor Plus 488 (ThermoFisher, cat#A32731) and goat anti-guinea pig IgG (H+L) Alexa Fluor 594 (ThermoFisher, cat#A11076) diluted to 1:1500. Coverslips were mounted using 20 µl antifade mountant (ThermoFisher, cat#P36971).

Images were captured using the Zeiss Axio Imager.Z2 Upright Fluorescence Microscope with ApoTome.2 (Fig 2A-C) or Olympus Fluoview FV1000 (Fig 2D-1, Fig 3A-I). The brightness, contrast, and colour balance of each image was adjusted using FIJI in ImageJ. Note that there are no SNb antibodies available for any species.

### Sample preparation for mass spectrometry analysis

On the afternoon before dissection, 30 males and 30 females were selected randomly and transferred into 30 breeding traps with one male and one female respectively separated by a transparent divider to prevent spawning. On the second day, these 30 couples were divided randomly in 3 groups of 10 couples each and anesthetized at 3 separate timepoints (Fig S4). Following anesthesia and spinal transection, brain, pituitary, and gonads were collected separately, placed in a homogenization buffer on ice and homogenized by ultrasonic homogenizer (Fisherbrand; Cat# FB120110) for 3-5 sec and processed for mass spectrometry as described previously (29). The homogenized samples were centrifuged at 4°C in 13,200 rpm for 10 min. After that, the supernatant was transferred and dehydrated in centrifugal vacuum concentrator (Labconco; Cat#7810014) at 45°C. The dehydrated supernatant samples were resuspended in loading buffer and subjected to SPE in a 96-well format. The purified samples were dehydrated under vacuum. Before loading into the LC-MS system, the samples were resuspended in loading buffer. All peptides were prepared by standard Fmoc/tBu chemistry and purified by HPLC (>95%) in our laboratory (29) and the steroids were from Sigma-Aldrich Inc. (Table S4). All other details are in the SI materials and methods.

### Ovulation induction

Seventy-two adult female zebrafish (7-8 mpf) were selected and isolated from males for 2 weeks. On the experimental day, they were randomly separated into 4 groups (n=18) and anaesthetized in 4.3% tricaine solution and injected i.p. with saline (pH=7; 10 µL/g body weight), SNa1-14 (1 nmol/g body weight) or SNa 1-34 (1 nmol/g body weight) using a 32-gauge Hamilton syringe. The Lh analog hCG (50 IU/g body weight) (Millipore, cat# 230734-1MG) served as a positive control for ovulation induction (27). At 6 h post injection, the fish were anesthetized on ice and ovaries were dissected. To weaken the attachment between oocytes and follicular layer, the dissected ovaries were incubated for 1 hour in modified Cortland’s medium without calcium and magnesium (68), and the numbers of ovulated and non-ovulated eggs for each fish were counted under a dissecting microscope.

### Effects of SNa on reproductive gene expression

Females were separated from males and kept in groups of 12 in 3L tanks for two weeks before treatments. On the experimental day, zebrafish (n=16) were randomly assigned to three groups and injected between 10:00-11:00 with saline, SNa1-14 (1 nmol/g body weight) or SNa 1-34 (1 nmol/g body weight). At 3 and 6 h after injection, 8 zebrafish in each group were sacrificed. Telencephalon, hypothalamus, pituitary, and ovaries were dissected from each individual and collected in 1.5 ml Eppendorf tubes, flash frozen in liquid nitrogen and kept at −80°C until RNA extraction. The total RNA isolation, cDNA synthesis, primer design (http://bisearch.enzim.hu/) and digital droplet PCR (ddPCR) followed our published methods (27). The annealing temperature of all the primers were 58°C. The primers (Integrated DNA Technologies) were validated by RT-PCR and the single product amplification was confirmed by the melting curve methods (Table S5). The PCR products were sequenced and confirmed using Basic Local Alignment Search Tool (BLAST).

### Data analysis

The LC-MS data is presented as mean+SEM (n=10). All the results were statistically analyzed on GraphPad Prism 9.0 software. The Kruskal-Wallis one-way analysis of variance on ranks followed by a Tukey test was performed to analyse time-dependent changes in hormone levels in each tissue. Non-linear regression analysis was performed to determine potential relationships between reproductive hormones and SNa peptides. The ddPCR data were analyzed using the Bio-Rad QuantaSoft 1.7 software. The PCR data were normalized using the Norma-Gene algorithm (69) and presented as mean+SEM (n=8). Normality and homoscedasticity were tested using the Shapiro-Wilk and Levene’s tests, respectively. Means were compared using two-way ANOVA followed by Tukey’s multiple comparisons test. The number of females ovulating in each treatment group was analyzed using Fisher’s Exact while % ovulated eggs was compared using the Mann-Whitney U test.

## Data Availability

All data, methods, and results of statistical analyses are reported in this paper and associated SI Appendix. We welcome any specific inquiries.

## ACKNOWLEDGMENTS

This research was funded by the China Scholarship Council (to D. Peng), the Natural Sciences and Engineering Research Council of Canada Discovery Grant (RGPIN-2021-03174 to V.L.T.), the University of Ottawa Research Chair program (to V.L.T.), the University of Ottawa International Research Acceleration Program (to V.L.T. and W.H.), the International Partnership Program of Chinese Academy of Sciences (Grant No 152342KYSB20180019 to W.H. and V.L.T.), St. Louis University (to D.Z.), and the Israel Science Foundation (to B.L.-S.). The authors acknowledge with appreciation the help of E. Ramsay for care of animals, and A. Ochalski and J. Liang for microscopy. The senior author recognizes that he works and lives on the unceded Algonquin Territory of the Three Fire Confederacy, Anishinaabewaki.

