## Supplemental files for "Hormonal dynamics reveal a stimulatory role for secretoneurin in zebrafish ovulation"

**This supporting PDF file includes:**
Supporting Information text
Table S1 to S5
Figs. S1 to S5
Supporting Information references

### Supporting tables and figures

| Peptide | | T1 | T2 | T3 |
|---|---|---|---|---|
| **Gnrh2** | Brain | 167.2±43.1 | 241.2±42.5 | 225.1±57.7 |
| | Pituitary | 3.6±1.7 | 0.5±0.0 | 5.7±2.9 |
| | Gonads | 23.4±16.1 | 3.8±2.3 | 19.5±14.1 |
| **Avp** | Brain | 47.5±23.6 | 75.4±26.1 | 90.9±32.9 |
| | Pituitary | 1638.0±433.3 | 1323.0±194.3 | 1374.0±288.3 |
| | Gonads | 267.4+93.9 | 453.2±301.8 | 409.7±255.6 |
| **Kiss1** | Brain | 128.8±46.6 | 57.5±24.1 | 23.2±8.1 |
| | Pituitary | 5.6±4.0 | 2.1±0.7 | 7.5±4.0 |
| | Gonads | 45.2±44.6 | 20.3±19.8 | 0.7±0.2 |
| **Kiss2** | Brain | 417.6±262.7 | 145.4±100.3 | 208.1±136.1 |
| | Pituitary | 870.2±264.9 | 727.7v246.6 | 567.8±237.3 |
| | Gonads | 646.0±226.6 | 580.7±239.9 | 184.3±149.5 |

**Table S1. Neuropeptide levels in the zebrafish periovulatory period**. The timepoints represent (T1) 02:00h, (T2) 05:00h and (T3) 08:30h. Results are presented as mean+SEM (n=10). None of these hormones exhibited periovulatory changes (p>0.05).

| Name | Amino acid sequence | | | | | | | | | | | | | | | | | | | | | | | | | | | | | | | | | |
|---|---|---|---|---|---|---|---|---|---|---|---|---|---|---|---|---|---|---|---|---|---|---|---|---|---|---|---|---|---|---|---|---|---|---|
| **SNa1-34** | **T** | **N** | **E** | **N** | **A** | **E** | **E** | **Q** | **Y** | **T** | **P** | **Q** | **K** | **L** | **A** | **T** | **L** | **Q** | **S** | **V** | **F** | **E** | **E** | **L** | **S** | **G** | **I** | **A** | **S** | **S** | **K** | **T** | **N** | **T** |
| SNa1-19 | T | N | E | N | A | E | E | Q | Y | T | P | Q | K | L | A | T | L | Q | S | | | | | | | | | | | | | | | |
| SNa1-18 | T | N | E | N | A | E | E | Q | Y | T | P | Q | K | L | A | T | L | Q | | | | | | | | | | | | | | | | |
| SNa1-14 | T | N | E | N | A | E | E | Q | Y | T | P | Q | K | L | | | | | | | | | | | | | | | | | | | | |
| SNa1-12 | T | N | E | N | A | E | E | Q | Y | T | P | Q | | | | | | | | | | | | | | | | | | | | | | |
| SNa1-9 | T | N | E | N | A | E | E | Q | Y | | | | | | | | | | | | | | | | | | | | | | | | | |
| **SNb1-31** | **A** | **T** | **E** | **D** | **L** | **D** | **E** | **Q** | **Y** | **T** | **P** | **Q** | **S** | **L** | **A** | **N** | **M** | **R** | **S** | **I** | **F** | **E** | **E** | **L** | **G** | **K** | **L** | **S** | **A** | **A** | **Q** | | | |
| SNb1-19 | A | T | E | D | L | D | E | Q | Y | T | P | Q | S | L | A | N | M | R | S | | | | | | | | | | | | | | | |
| SNb1-18 | A | T | E | D | L | D | E | Q | Y | T | P | Q | S | L | A | N | M | R | | | | | | | | | | | | | | | | |
| SNb1-18* | A | T | E | D | L | D | E | Q | Y | T | P | Q | S | L | A | N | *M* | R | | | | | | | | | | | | | | | | |
| SNb1-17 | A | T | E | D | L | D | E | Q | Y | T | P | Q | S | L | A | N | M | | | | | | | | | | | | | | | | | |
| SNb1-17* | A | T | E | D | L | D | E | Q | Y | T | P | Q | S | L | A | N | *M* | | | | | | | | | | | | | | | | | |
| SNb1-16 | A | T | E | D | L | D | E | Q | Y | T | P | Q | S | L | A | N | | | | | | | | | | | | | | | | | | |
| SNb1-15 | A | T | E | D | L | D | E | Q | Y | T | P | Q | S | L | A | | | | | | | | | | | | | | | | | | | |
| SNb1-14 | A | T | E | D | L | D | E | Q | Y | T | P | Q | S | L | | | | | | | | | | | | | | | | | | | | |
| SNb1-13 | A | T | E | D | L | D | E | Q | Y | T | P | Q | S | | | | | | | | | | | | | | | | | | | | | |
| SNb1-12 | A | T | E | D | L | D | E | Q | Y | T | P | Q | | | | | | | | | | | | | | | | | | | | | | |
| SNb1-11 | A | T | E | D | L | D | E | Q | Y | T | P | | | | | | | | | | | | | | | | | | | | | | | |
| SNb1-10 | A | T | E | D | L | D | E | Q | Y | T | | | | | | | | | | | | | | | | | | | | | | | | |
| SNb1-9 | A | T | E | D | L | D | E | Q | Y | | | | | | | | | | | | | | | | | | | | | | | | | |
| SNb12-31 | | | | | | | | | | | | Q | S | L | A | N | M | R | S | I | F | E | E | L | G | K | L | S | A | A | Q | | | |
| SNb19-31 | | | | | | | | | | | | | | | | | | | S | I | F | E | E | L | G | K | L | S | A | A | Q | | | |
| SNb20-31 | | | | | | | | | | | | | | | | | | | | I | F | E | E | L | G | K | L | S | A | A | Q | | | |
| SNb19-30 | | | | | | | | | | | | | | | | | | | S | I | F | E | E | L | G | K | L | S | A | A | | | | |
| SNb19-29 | | | | | | | | | | | | | | | | | | | S | I | F | E | E | L | G | K | L | S | A | | | | | |
| SNb19-28 | | | | | | | | | | | | | | | | | | | S | I | F | E | E | L | G | K | L | S | | | | | | |

**Table S2.** Identified secretoneurin fragments from zebrafish brain and pituitary by untargeted peptidomics. *Note than SNb1-18 and SNb1-7 also exist with both non-oxidated and oxidated methionine as indicated by *M*.

| REGRESSION ($R^2$) | | SNa1-34 | SNa1-14 |
|---|---|---|---|
| Brain | Gnrh2 | 0.4710 § | **0.5765** ¶* |
| | Gnrh3 | **0.7067** ¶** | 0.1124 |
| | Oxt | 0.3133 | 0.2629 ¶ |
| | Avp | 0.0626 | 0.1391 |
| | Kiss1 | 0.1346 | 0.0692 |
| | Kiss2 | 0.0526 | 0.0518 |
| | E1 | 0.1312 | 0.0169 |
| | E2 | 0.3809 § | 0.0045 |
| | E3 | 0.0495 | 0.0692 |
| | T | 0.0994 | 0.1207 |
| | 11-KT | 0.1363 | 0.1835 |
| Pituitary | Gnrh2 | 0.0292 | 0.2058 |
| | Gnrh3 | 0.0565 | 0.3824 § |
| | Oxt | 0.1979 | 0.0940 |
| | Avp | 0.1613 § | 0.0678 |
| | Kiss1 | 0.3911 | 0.1519 |
| | Kiss2 | 0.0742 | 0.2659 ¶ |
| | E1 | 0.1795 | 0.4147 |
| | E2 | 0.1472 § | 0.1026 |
| | E3 | 0.3671 | 0.0390 |
| | T | 0.0404 | 0.0692 |
| | 11-KT | 0.0217 | 0.0215 |
| Ovary | Gnrh2 | 0.0158 | 0.1039 |
| | Gnrh3 | 0.1220 | 0.0422 |
| | Oxt | 0.0814 | 0.0321 ¶ |
| | Avp | 0.1878 | 0.0394 |
| | Kiss1 | 0.0322 | 0.4401 § |
| | Kiss2 | 0.1301 | 0.1832 |
| | E1 | 0.2565 | 0.0386 |
| | E2 | 0.1319 | 0.1014 |
| | E3 | 0.1458 | 0.0812 |
| | T | 0.0492 | 0.2075 |
| | 11-KT | 0.1958 | 0.2380 |

**Table S3. Regression analysis of reproductive hormones and SNa related peptides in the periovulatory period in zebrafish brain.** Results are presented as $R^2$ value to show the goodness of fit (n=30). * $0.5 < R^2 < 0.7$, ** $R^2 > 0.7$. The § indicates a third-order polynomial; the ¶ indicates a fourth-order polynomial. All other $R^2$ are not significant (<0.50) and are second-order polynomials. The best-fit models for non-linear regression were determined using GraphPad Prism 9.0 software. The software compares the third-order polynomial as the null hypothesis with the fourth-order polynomial as the alternative hypothesis. It accepts the alternative hypothesis when $p<0.05$ while it accepts the null hypothesis when $p>0.05$. If $R^2 < 0.3$, it is considered as not related; if $0.3 < R^2 < 0.5$, it is considered as weakly related; if $0.5 < R^2 < 0.7$, it is considered as moderately related; if $R^2$ value > 0.7, it is considered as strongly related (1).

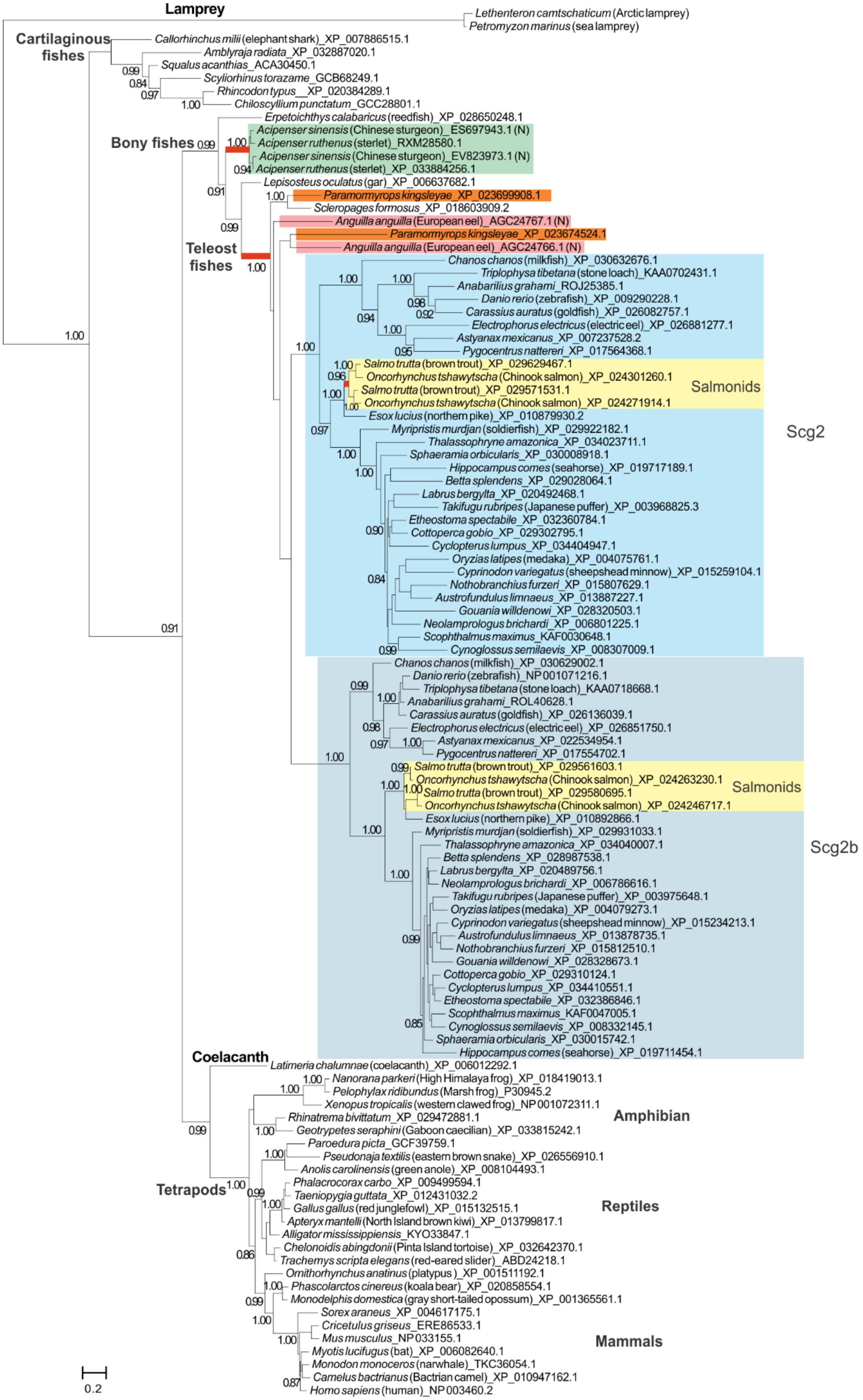


**Fig S1. Phylogenetic analysis of Scg2a and Scg2 homologs from representative jawed vertebrates**. The tree was generated using the FastTree program with the major branches supported by the SH-like local support values. Each sequence is represented by the species name followed by the NCBI accession ID number. The genome duplication events found in *Acipenser ruthenus*, ancestral to teleost fishes, and *Salmo trutta* are indicated by thick red branches and their paralogous groups are highlighted in different colored backgrounds.

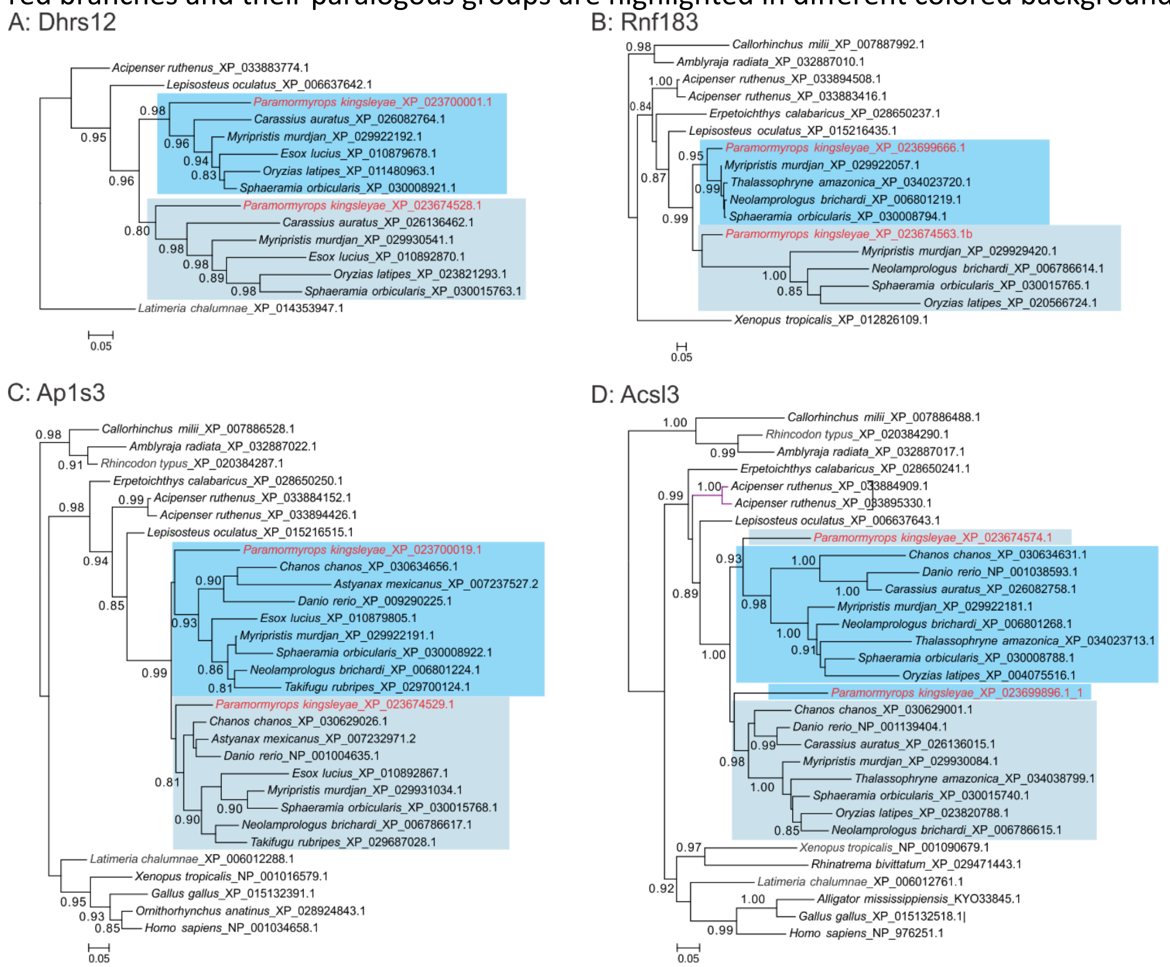


**Fig S2. Phylogenetic analysis of four neighboring genes of the Scg2a and Scg2b containing loci.** Shown are (A) Dhrs12, (B) Rnf183, (C) Ap1s and (D) Acsl3. The tree was generated using the FastTree program with the major branches supported by the SH-like local support values. Each sequence is represented by the species name followed by the NCBI accession ID number. The two paralogous groups after the genome duplication are highlighted in different background colors.

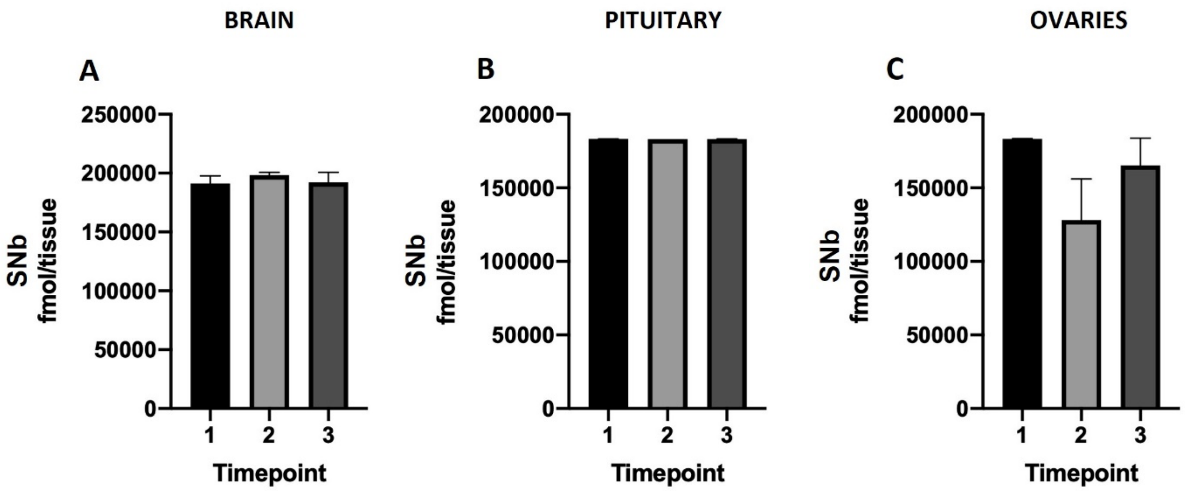


**Fig S3. SNb levels during the periovulatory period in brain, pituitary and ovaries of female zebrafish.** The timepoints represent (1) 02:00h, (2) 05:00h and (3) 08:30h. SNb1-31 in brain (A), pituitary (B) and ovaries (C).

Results are presented as mean+SEM (n=10). Note the different scales on the Y-axes. The Kruskal-Wallis one-way analysis of variance on ranks was performed followed by a Tukey test but revealed no changes (p>0.05).

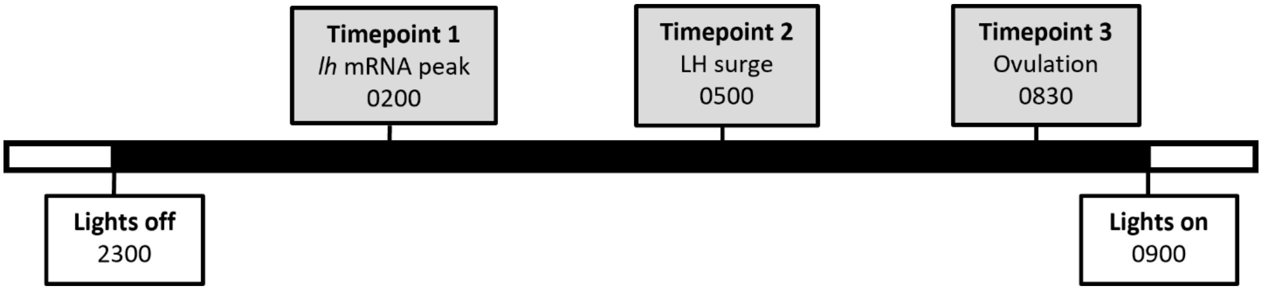


**Fig S4.** Timepoints of zebrafish dissections in the periovulatory period

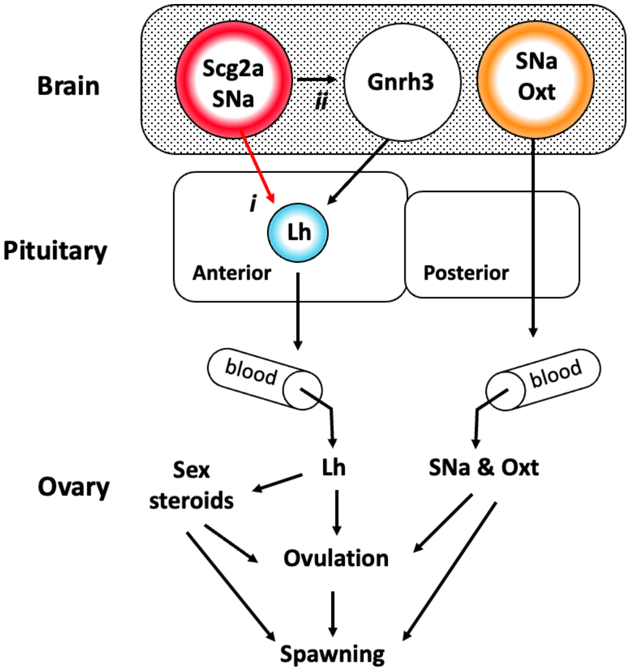


**Fig S5.** Summary of key findings and working model for the role of SNa1-34 in female zebrafish reproduction. The Scg2a precursor protein is selectively processed by prohormone convertases to the bioactive peptide SNa1-34. Immunocytochemistry revealed SNa and OXT are colocalized in preoptic neurons. We found that SNa+OXT neurons project to the neurointermediate lobe in the posterior pituitary. While direct evidence is still scant, localization of SNa and OXT in nerve terminals implies release into the circulation, where the peptides may influence ovulation and spawning. Neuronal fibres immunoreactive for SNa but OXT negative entered the proximal pars distalis (anterior pituitary) and terminated in close proximity to Lh gonadotrophs in Tg(*lhb*-RFP X *fshb*-eGFP) females. Regression analysis revealed that SNa1-34 (shown as SNa on the figure) and Gnrh3 peptide levels in the brain increase concomitantly at the time of the Lh surge. In the pituitary, SNa1-34 peptide levels are highest prior to ovulation. Experimental data demonstrated that 3 hr following i.p. injection of synthetic SNa1-34 increases mRNA levels for *gnrh3* in the hypothalamus, *lhb* and *cga* in the pituitary, and *lhcgr* and *npr* in the ovary. These are key indicators of activation of genes at each level of the HPG axis that led to ovulation 6 hr after a single SNa1-34 injection. SNa1-34 can enhance Lh cell activity through a neuroendocrine pathway (i) and by stimulation of Gnrh3 neurons (ii). This model is also supported by previous observations in mutant lines (2) where it was demonstrated that hypothalamic *gnrh3*, and pituitary *lhb* and *cga* are lower in *scg2a-/-* females compared to wildtype. In the same mutant line, SNa1-34 injection in the subfertile *scg2a-/-; scg2b-/-* double mutant zebrafish enhanced spawning by ~3 fold, like the effects of the Lh analog hCG (2).

### Supporting Information (SI) Text

### SI Materials and Methods

**Sequence searches and phylogenetics analysis.** To comprehensively collect the homologous sequences of Scg2, iterative profile searches were performed by using PSI-BLAST (3) against non-redundant (nr) protein database within the National Center of Biotechnology Information (NCBI) with an expected value (e-value) of 0.005. We also utilized the TBLASTN program to identify the unannotated Scg2 homologs from sequenced genomes of several fish species, such as *Acipenser sinensis* (Chinese sturgeon) and *Anguilla anguilla* (European eel), several basal vertebrates, such as *Petromyzon marinus* (sea lamprey), *Lethenteron camtschaticum* (Arctic lamprey), and the protochordate *Ciona intestinalis* which is the outside of vertebrate clade. The Scg2 homologs from several representative vertebrate species were collected and multiple sequence alignments were promoted by using the KALIGN (4). An unrooted phylogenetic tree was reconstructed using an approximately-maximum-likelihood method implemented in the FastTree 2.1 program with the JTT evolutionary model and the discrete gamma model with 20 rate categories (5). The tree was rendered using MEGA Tree Explorer (6).

**Tissue processing for immunocytochemistry**. After washing in PBS 15 min for three times, the decalcified samples from wildtype females were dehydrated in ethanol (EtOH) in a series of 30-minute incubations in increasing concentrations of EtOH (30, 50, 70, 80, 99%). Next, samples were incubated in the xylene for 2 X 30 min and transferred into melted paraffin at 60°C for 1 hour, then incubated in clean paraffin at 60°C overnight. The next morning, samples were embedded in metal molds and prepared for sectioning (8-10 µm) on the Microm HM350 (Heidelberg) or Leica RM2255 microtomes and mounted on microscope glass slides (Fisherbrand, cat#12-550-15). One the day of immunofluorescence labelling, the sections were incubated in Coplin jars with xylene 2 x 20 min and rehydrated in EtOH with a series of 2 X 10-minute incubations in decreasing concentrations of EtOH (99, 95, 70%) and finally in PBS. For cryosectioning (20 µm) of Tg(*lhb*-RFP X *fshb*-eGFP) females, samples were incubated for 6 hrs in increasing concentrations of sucrose (10, 20, 30%) and embedded in plastic molds with O.C.T compound (Fisher Science, cat# 23-730-571) and frozen at -80°C. Cryosectioning was performed on the Leica CM1850 Cryostat when operating temperature at -18°C and chamber temperature at -20°C. The cryosections were mounted on microscope glass slides (Fisherbrand, cat#12-550-15) and stored at -80°C until use. For immunofluorescence labelling, slides were thawed at room temperature and washed in 2 X 10 minutes PBS. In Coplin jars, slides were incubated in 0.01M sodium citrate solution at 80°C for 30 minutes, and then cooled to room temperature for antigen retrieval. Slides were incubated in blocking buffer with 1% skim milk and 0.3% Triton-100 for 30 minutes prior to incubation with primary and secondary antibodies.

**Mass spectroscopy**. All tissues were dissected and placed directly into 1.5 mL centrifuge tubes with homogenization buffer (90% MeOH, 9% water, and 1% acetic acid v/v/v) and processed following our established methods (7). A probe homogenizer (Fisherbrand, Cat#FB120110) was operated at 10 Watts for 20 secs. The pellet and supernatant of the homogenate were evaporated in a speed-vac after centrifugation. Dried pellets were solubilized in 200 µL 8 M urea, 50 mM ammonium bicarbonate (pH 8.0). Dithiothreitol was added to a final concentration of 10 mM and incubated at room temperature for 1hr for reduction. Iodoacetamide was added to a final concentration of 20 mM and incubated at room temperature for 40 min in dark for alkylation. The samples were diluted 5-fold in 50 mM ammonium bicarbonate and digested by 1 µg trypsin/20 µg sample protein overnight at room temperature. The SPE plates for both peptide-steroid co-extraction and the protein hydrolysate extraction were prepared by packing extraction sorbent (Dr. Maisch, r10. aq) into filtered 96-well plates (OROCHEM, OF1100-7PE). Recovery rates of standards for SPE peptide-steroid co-extraction was >85% for 0.01-10 nmol loading.

Peptides were assembled in-lab on the Intavis Multipeptide RSi peptide synthesis machine (Köln, Germany) using standard Fmoc peptide synthesis chemistry as described (7). Peptide purification (>95%) was performed on a 4.6 mm × 150 mm Phenomenex Luna Omega 5µm C18 column (CA, USA) an Agilent 1100 HPLC system. The purity of the products was verified by nano flow UHPLC-MS/MS using 75µm × 100 mm analytical column packed in-house with reverse phase Magic C18AQ resins (1.9µm; 120-Å pore size; Dr. Maisch GmbH, Ammerbuch, Germany), confirming that all samples were free of significant chiral isomer contamination. See Table S5 for standards.

Untargeted peptidomic analysis (7) was used to identify the SNa and SNb fragmental peptides presented in Table S2. Targeted peptidomic analysis was performed to confirm the predicted amino acid sequence of SNa1-34 and SNb1-31 peptides in comparison to standard compounds. To confirm the peptide sequences, we identified 5 of the most intense product ions (MS2) of the SNa and SNb peptides in the analyte samples for the same parent ion (MS1) at the same retention time with 30s detection window. All peptide and steroid MS analyses were done by HPLC-ESI-MS/MS (7). Peptide structures were confirmed by three MS2 ions and steroid structures were confirmed by one MS2 ions. Chromatograms were plotted by accurate MS1 with 10 ppm mass tolerance and 120 sec identification windows. When there were over 5 data points to form a chromatographic peak by Genesis peak detection, the level was determined by comparing peak area to standard curves. The system consisted of an Agilent 1100 micro-HPLC system (California, USA) coupled with an LTQ-Orbitrap mass spectrometer (California, USA) equipped with a nano-electrospray interface. The mobile phases consisted of 0.1%(v/v) FA in water as buffer A and 0.1% (v/v) FA in acetonitrile as buffer B. Peptide separation was performed on a 75µm × 100 mm analytical column packed in-house with reverse phase Magic C18AQ resins (1.9µm; 120-Å pore size; Dr. Maisch GmbH, Ammerbuch, Germany). Briefly, the sample was loaded on the column using 98% buffer A at a flow rate of 1.5µL/min for 15min. Then, a gradient from 5% to 35% buffer B (20~50% for APols depletion test) was performed in 120 min at a flow rate of ~300nL/min obtained from splitting a 20 µL/min through a restrictor. The MS method consisted of one full MS scan from 300 to 1700 m/z followed by data-dependent MS/MS scan of the 5 most intense ions, with dynamic exclusion repeat count of 2, and repeat duration of 90 s. As well, for the experiments on the Orbitrap MS the full MS was in performed in the Orbitrap analyzer with R = 60,000 defined at m/z 400, while the MS/MS analysis were performed in the LTQ. To improve the mass accuracy, all the measurements in Orbitrap mass analyzer were performed with internal recalibration ("Lock Mass"). On the Orbitrap, the charge state rejection function was enabled, with single and "unassigned" charge states rejected.

The raw files generated by the LTQ-Orbitrap were processed and analyzed using MaxQuant, Version 1.2.2.5 using the Uniprot protein FASTA database including commonly observed contaminants. The following parameters were used: cysteine carbamidomethylation was disabled; methionine oxidation, protein N-terminal acetylation, and enzyme digestion disabled. Precursor ion mass tolerances were 7 ppm, and fragment ion mass tolerance was 0.8 Da for MS/MS spectra. The false discovery rate (FDR) for peptide and protein was set at 1% and a minimum length of six amino acids was used for peptides identification. Quantification was performed using normalized LFQ intensity.

| Standard compound | Amino acid sequence or other information |
|---|---|
| Oxt (Isotocin) | H-CYISNCPIG-CONH2 [Disulfide C1-C6] |
| Avp (Vasotocin) | H-CYIQNCPRG-CONH2 [Disulfide C1-C6] |
| Kisspeptin 1 (Kiss1) | H-Y(PO3)NLNSFGLRY-CONH2 |
| Kisspeptin 2 (Kiss2) | H-FNYNPFGLRF-CONH2 |
| Gnrh2 | Pyr-HWSHGWYPG-CONH2 |
| Gnrh3 | H-Pyr-HWSYGWLPG-CONH2 |
| SNa1-34 | H-TNENAEEQYTPQKLATLQSVFEELSGIASSKTNT-CONH2 |
| SNa1-14 | H-TNENAEEQYTPQKL-CONH2 |
| SNa1-18 | H-TNENAEEQYTPQKLATLQ-CONH2 |
| SNa19-34 | H-SVFEELSGIASSKTNT-CONH2 |
| SNb1-31 | H-ATEDLDEQYTPQSLANMRSIFEELGKLSAAQ-CONH2 |
| SNb1-17 | H-ATEDLDEQYTPQSLANM-CONH2 |
| SNb19-31 | H-SIFEELGKLSAAQ-CONH2 |
| Estrone (E1) | Sigma-Aldrich; Cat. # E9750 |
| Estradiol (E2) | Sigma-Aldrich; Cat. # E1024 |
| Estriol (E3) | Sigma-Aldrich; Cat. # E1253 |
| Testosterone (T) | Sigma-Aldrich; Cat. # Y0002174 |
| 11-Ketotestosterone (11-KT) | Sigma-Aldrich; Cat. # K8250 |

**Table S4.** Standard compounds used for hormone analysis by LC-MS

| Gene (abbreviation) | Tissue analysed | Sequence (5`-3`) | Amplicon size (bp) |
|---|---|---|---|
| Gonadotropin-releasing hormone 2 (*gnrh2*) | Tel, Hyp | Fw: CAGAGGTTTCAGAGGAAGTGAAGC<br>Rv: TGAGGGCATCCAGCAGTATTG | 154 |
| Gonadotropin-releasing hormone 3 (*gnrh3*) | Tel, Hyp | Fw: ATGGAGGCAACATTCAGGATGT<br>Rv: CCTTTCAGAGGCAAACCTTCA | 131 |
| Isotocin (*oxt*) | Tel, Hyp | Fw: GATCTGCTGCTGAAGCTCCT<br>Rv: TACAAAAGTGGGTGGCGAGT | 134 |
| Vasotocin (*avp*) | Tel, Hyp | Fw: AGGTCTGCATGGAAGAGGAG<br>Rv: CTGCCTTCAGGACAGTCTGG | 146 |
| Follicle stimulating hormone beta (*fshb*) | Pituitary | Fw: TGTGGAGAGCGAAGAATGTG<br>Rv: AGACCTTCTGGGTGTGCTGT | 116 |
| Gonadotropin- releasing hormone receptor 2 (*gnrhr2*) | Pituitary | Fw: TCCTCAACCCTCTGTCCATC<br>Rv: TGCTTTGGGGAATCAATCTC | 123 |
| Glycoprotein alpha (*cga*) | Pituitary | Fw: CTGCTGCTTTTCGAGAGCTT<br>Rv: AGTGGCAGTCTGTGTGGTTG | 155 |
| Luteinizing hormone beta (*lhb*) | Pituitary | Fw: AATGCCTGGTGTTTCAGACC<br>Rv: AACAGTCGGGCAGGTTAATG | 144 |
| *cyp19a1a* | Ovary | Fw: ACTGAAAGGGCTCAGGACAA<br>Rv: TCCAACTGACCTGGAATGTG | 174 |
| Luteinizing hormone receptor (*lhcgr*) | Ovary | Fw: TGAAAGAGCAGCCAGGTACT<br>Rv: TGCTAAATTTCCTTTCGCCG | 166 |
| Nuclear progesterone receptor (*npr*) | Ovary | Fw: GGGCCACTCATGTCTCGTCTA<br>Rv: TCTCCACTCTGAAAATATGTGGACTTT | 198 |
| Steroidogenic acute regulatory protein 1 (*star1*) | Ovary | Fw: GGTCTGAGGAAGAATGCAATGAT<br>Rv: CCAGGTCCGGAGAGCTTGT | 169 |

**Table S5.** Forward (Fw) and reverse (Rv) primers for ddPCR with sequence-verified amplicon sizes.
